# MAIT cells exacerbate liver fibrosis by downsizing the intrahepatic regulatory T cell compartment

**DOI:** 10.64898/2026.04.15.718691

**Authors:** Nicole I. Wang, Valeryia Shydlouskaya, Katelyn R. Reid, Agetha Mahendran, Angela Schincaglia, Brian A. Keller, Shereen Q. Zia, Ahmad R. Movasseghi, Julius Haruna, Dale I. Godfrey, S.M. Mansour Haeryfar

**Affiliations:** Department of Microbiology and Immunology, Western University, London, ON, Canada; Collaborative Specialization in Machine Learning in Health and Biomedical Sciences, Western University, London, ON, Canada; Department of Pathology and Laboratory Medicine, Schulich School of Medicine & Dentistry, Western University, London, ON, Canada; Department of Pathology and Laboratory Medicine, London Health Sciences Centre, London, ON, Canada; Charles River Laboratories, Laval, QC, Canada; Department of Microbiology and Immunology, The Peter Doherty Institute for Infection and Immunity at the University of Melbourne, Melbourne, Victoria, Australia; Department of Oncology, Western University, London, ON, Canada; Division of General Surgery, Department of Surgery, Western University, London, ON, Canada; Division of Clinical Immunology and Allergy, Department of Medicine, Western University, London, ON, Canada; Lawson Health Research Institute, London Health Sciences Centre, London, ON, Canada

**Author notes:** **Corresponding Author:** Dr. S.M. Mansour Haeryfar, Department of Microbiology and Immunology, Western University, 1151 Richmond Street, London, Ontario, N6A 5C1, Canada.

**Keywords:** MAIT cells, liver fibrosis, tissue repair, type-17 immunity, regulatory T cells

## Abstract

Mucosa-associated invariant T (MAIT) cells have been paradoxically implicated in both tissue repair and fibrosis. However, when and how they modulate fibrogenesis in the injured liver remain unclear. Here, using the carbon tetrachloride-induced model of liver injury in MR1- and MAIT cell-sufficient and -deficient mice, we identify MAIT cells as an early driver of fibrogenesis. The presence of MAIT cells exacerbated hepatocellular injury, myofibroblast activation, and matrix deposition early in the course of fibrosis development, but not at later stages. This was accompanied by rapid polarization of hepatic MAIT cells toward a MAIT17 phenotype and enrichment of pro-fibrotic transcriptional programs. Concurrently, MAIT cells acquired an exhaustion-associated phenotype while still retaining their effector functions. Mechanistically, we demonstrate that MAIT cells limit hepatic regulatory T (T_reg_) cell accumulation, accompanied by reduced Ki-67 and CXCR3 levels in the latter population, suggesting their impaired proliferation and tissue recruitment. Furthermore, T_reg_ cell inactivation reversed MAIT cell-dependent differences in the severity of fibrosis, establishing T_reg_ cells as a key downstream mediator. Together, these findings identify MAIT cells as early orchestrators of fibrogenesis and reveal a novel MAIT-T_reg_ axis that can be considered a potential therapeutic target in the early stages of fibrotic diseases.

## INTRODUCTION

Advanced liver fibrosis affects an estimated 3.3% of the global population and contributes to approximately two million annual deaths associated with liver disease^1,2^. Pathological fibrosis arises when repeated hepatic injury from viral, metabolic, toxic or autoimmune aetiologies disrupts wound-healing responses and drives progressive scar formation^3,4^. The fibrogenic remodeling process is characterized by excessive accumulation of extracellular matrix components, primarily collagen, and distortion of normal liver architecture^3,4^. These structural changes are sustained by dynamic crosstalk between injured parenchymal cells, collagen-producing stromal cells such as hepatic stellate cells (HSCs), and diverse resident and recruited immune cell populations that regulate hepatic inflammation and repair^4^.

Mucosa-associated invariant T (MAIT) cells are a subset of innate-like T lymphocytes that comprise up to 50% of all T cells in the human liver^5^. The MAIT cell T cell receptor (TCR) is highly conserved and consists of an invariant α chain (typically Vα19-Jα33 in mice and Vα7.2-Jα33 in humans) paired with a restricted set of Vβ chains^6^. Unlike conventional T (T_conv_) cells that recognize peptide antigens, MAIT cells respond to riboflavin-derived microbial metabolites presented by monomorphic MHC-related protein 1 (MR1) molecules^7^. MAIT cells can also be activated in the absence of cognate antigen by pro-inflammatory cytokines, including interleukin (IL)-12 and IL-18^8^. Upon activation, MAIT cells rapidly secrete large quantities of inflammatory cytokines such as tumor necrosis factor (TNF), interferon (IFN)-γ and IL-17A, which enable them to regulate the function of various other cell types participating in immune and inflammatory responses^9^.

While the role of T_conv_ cells in fibrosis is well established^10^, the contribution of MAIT cells to tissue repair and fibrogenesis is only beginning to be elucidated. In acute models of tissue injury, MAIT cell activation has been linked to transcriptional tissue repair signatures and accelerated wound closure, suggesting a protective role^11–13^. In chronic liver disease, MAIT cells appear to be pathogenic as intrahepatic MAIT cells from patients with fibrosis exhibit features of chronic activation, exhaustion, and pro-fibrotic functionality^14,15^. Consistent with these observations, several studies on mouse models of chronic liver injury have suggested that MAIT cells worsen fibrosis^14,16^. In contrast, Jiang *et al.*^17^ reported that MAIT cells reduce the severity of fibrosis by enhancing natural killer (NK) cell-mediated cytotoxicity against HSCs. Therefore, although MAIT cells have been linked to both tissue repair and hepatic fibrosis, their overall role in fibrosis development in the injured liver remains ill-defined. In particular, the temporal dynamics of MAIT cell influence on fibrogenesis have not been explored. Also importantly, the mechanisms by which their effector programs shape the hepatic immune milieu *in vivo* are poorly understood.

In this study, we have employed the carbon tetrachloride (CCl_4_) model of liver injury and fibrosis and several MAIT cell-enriched and -deficient mouse strains to assess hepatic injury, fibrogenic remodeling, and immune cell infiltrates over time. We show that intrahepatic MAIT cells accelerate the onset and early progression of hepatic fibrosis, while their impact becomes less pronounced as fibrosis gains traction. We further demonstrate that hepatic MAIT cells acquire a robust pro-fibrotic type-17 (MAIT17) phenotype and that they reduce regulatory T (T_reg_) cell accumulation in the injured liver. Ultimately, these findings pinpoint an early window in which MAIT cells drive fibrogenesis and identify a previously unrecognized MAIT cell-T_reg_ cell axis that shapes a pro-fibrotic microenvironment.

## RESULTS

### MAIT cells promote hepatic damage and matrix accumulation at the onset of chronic injury

To explore the role of MAIT cells in liver damage and fibrosis, we administered CCl_4_ to MAIT cell-enriched *Mr1^+/+^* C57BL/6 (B6)-MAIT^CAST^ mice and their MAIT cell-deficient *Mr1^-/-^*B6-MAIT^CAST^ counterparts, which will hereafter be referred to as *Mr1^+/+^* and *Mr1^-/-^* mice, respectively. *Mr1^+/+^* mice exhibit elevated MAIT cell frequencies compared to the conventional wildtype B6 strain^18–20^, thus simulating MAIT cell numbers in human tissues including the liver. To induce chronic injury and fibrosis, mice received intraperitoneal (*i.p.*) CCl_4_ injections twice weekly, while control animals received a corn oil vehicle^14,21,22^ **(Fig. 1a)**. Animals were analyzed at either day 12 or day 26, representing the early and established stages of liver fibrosis, respectively^21^ **(Fig. 1a)**.

**Fig. 1.**
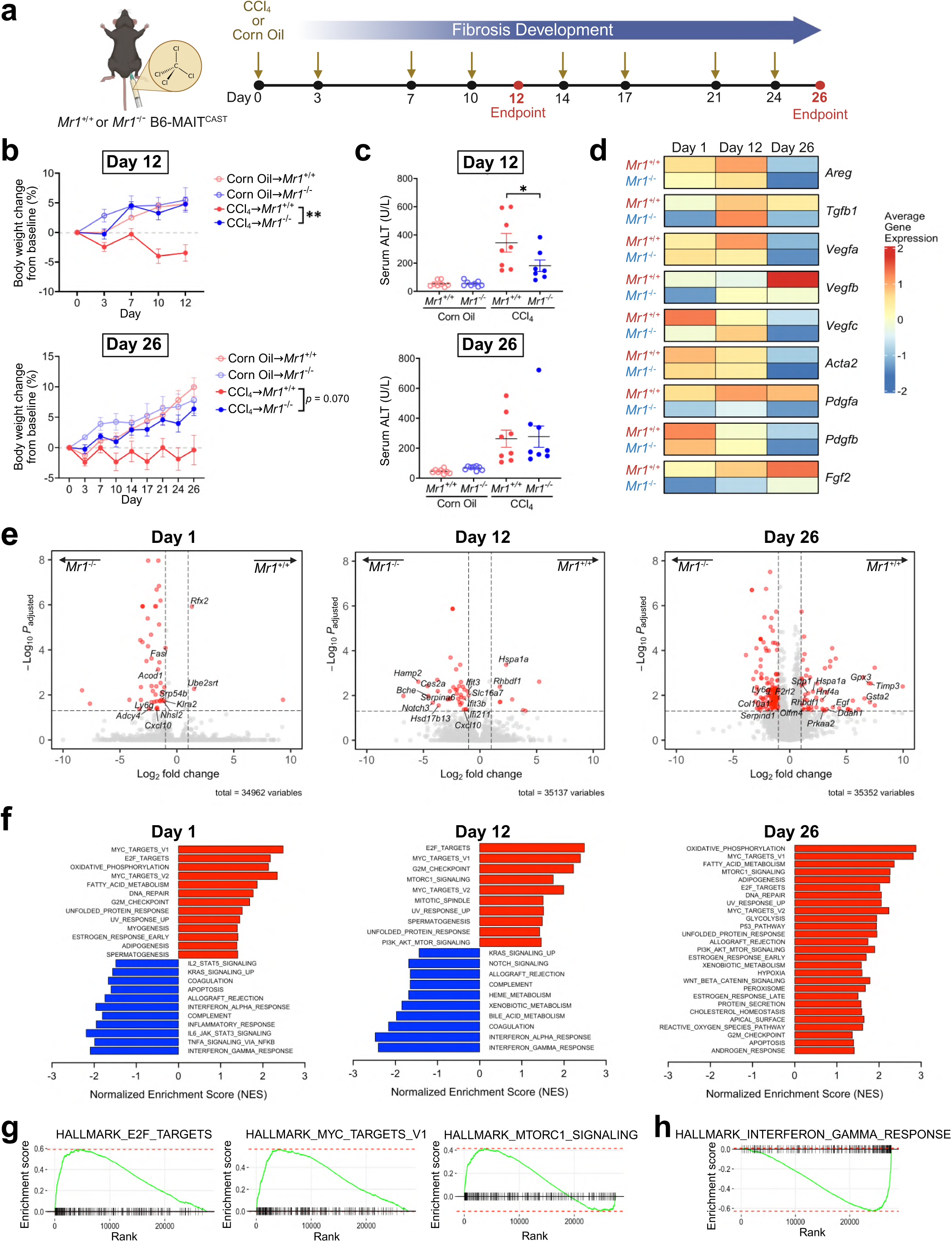
MAIT cells promote early hepatic injury and pro-fibrotic transcriptional programs. *Mr1^+/+^* or *Mr1^-/-^* mice were injected with CCl_4_ or with corn oil *i.p.* twice weekly until the indicated endpoints (a). Percent body weight changes from baseline were recorded over time for 12 or 26 days (b), and serum ALT levels were quantified by ELISA (c). Non-parenchymal HMNCs were isolated from CCl_4_-treated *Mr1^+/+^* and *Mr1^-/-^*mice on days 1, 12, and 26, and subjected to bulk RNA-seq. Mean expression of fibrosis-related genes is depicted in a heatmap (d). Differential gene expression between CCl_4_-treated *Mr1^+/+^* and *Mr1^-/-^* mice at days 1, 12, and 26 is presented as volcano plots (e). GSEA of differentially expressed genes identified significantly enriched Hallmark pathways, shown as normalized enrichment scores (NES) (f), with representative enrichment plots for selected pathways upregulated (g) and downregulated (h) in *Mr1^+/+^* mice relative to *Mr1^-/-^* mice, shown for Day 12. Data in (b) and (c) are pooled from 4 independent experiments. Data in (b-c) are presented as mean ± SEM. Statistical analyses were performed using two-way repeated measures ANOVA for (b) and two-way ANOVA for (c). * and ** denote differences with *p* < 0.05 and 0.01, respectively.

Differences associated with MAIT cells’ presence or absence became apparent early during liver injury. By day 12, CCl_4_-treated *Mr1^+/+^* mice exhibited greater body weight loss compared to *Mr1^-/-^* animals that maintained growth patterns similar to corn oil-treated controls **(Fig. 1b)**. A similar, but non-significant, trend was observed when mice were assessed on day 26 **(Fig. 1b)**. Moreover, serum alanine transaminase (ALT), a sensitive indicator of hepatocellular injury, was markedly elevated in CCl_4_-treated *Mr1^+/+^* mice on day 12, while no strain differences were observed on day 26 **(Fig. 1c)**. These findings suggest that MAIT cells potentiate early liver damage in the aftermath of CCl_4_ administration.

We next asked whether the presence or absence of MAIT cells was associated with distinct transcriptional programs in response to CCl_4_ over time. To this end, we performed bulk RNA sequencing (RNA-seq) on non-parenchymal hepatic mononuclear cells (HMNCs) isolated 1, 12 and 26 days after the initial CCl_4_ administration, with day-1 samples being included to capture acute injury preceding fibrotic remodeling. Fibrosis-related genes were consistently enriched in *Mr1^+/+^* mice compared to *Mr1^-/-^* animals across all time points examined, including genes involved in tissue repair (*Areg*, *Tgfb1*), angiogenesis (*Vegfa, Vegfb, Vegfc)*, and extracellular matrix regulation (*Acta2, Pdgfa, Pdgfb, Fgf2*) **(Fig. 1d)**. In addition, differential gene expression analyses revealed significantly elevated *Hspa1a* and *Rhbdf1*, genes associated with fibrosis progression^23,24^, in CCl_4_-treated *Mr1^+/+^* mice on both days 12 and 26 **(Fig. 1e)**. Additionally, gene set enrichment analysis (GSEA) demonstrated sustained enrichment of proliferation-related hallmark pathways in *Mr1^+/+^* mice, including E2F and MYC targets across all time points (days 1, 12, and 26), and mTORC1 signalling at later time points (days 12 and 26), all of which are known to promote fibrotic remodeling^25–28^ **(Fig. 1f-g)**. In contrast, the IFN-γ response pathway, which has been implicated in limiting fibrosis^29,30^, was downregulated in *Mr1*^+/+^ mice at early time points (days 1 and 12) **(Fig. 1f, h)**.

Histopathological analyses corroborated our transcriptomic findings. Accordingly, immunohistochemical staining for α-smooth muscle actin (α-SMA), a marker of activated myofibroblasts, revealed greater α-SMA^+^ areas in *Mr1^+/+^* livers compared with *Mr1^-/-^* mice on day 12, while no strain-dependent differences were evident on day 26 **(Fig. 2a)**. Similarly, collagen deposition, which was assessed by Picrosirius Red staining, was augmented in *Mr1^+/+^*mice on day 12, but not on day 26 **(Fig. 2b)**. These differences were observed in all four lobes of the livers **(Supplementary Fig. 1a-b)** and in both sexes **(Supplementary Fig. 1c-f)**. Blinded scoring by board-certified pathologists confirmed these findings, with higher Ishak fibrosis scores in *Mr1^+/+^* mice on day 12 **(Fig. 2c)**. No differences were detectable in ballooning or inflammation scores **(Supplementary Fig. 2a-b)**.

**Fig. 2.**
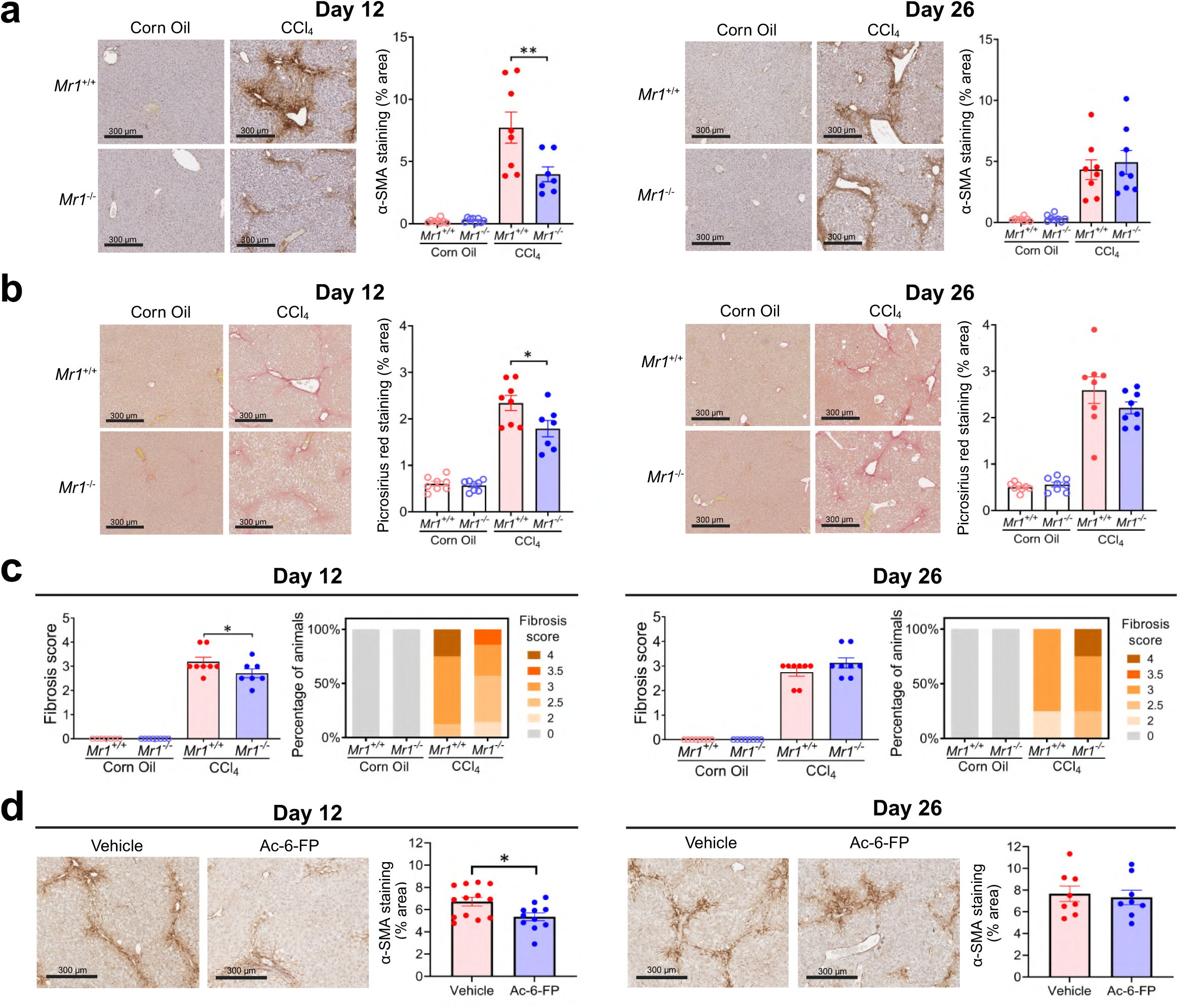
MAIT cells enhance early fibrogenesis in the liver. Formalin-fixed, paraffin-embedded liver sections from corn oil- or CCl_4_-treated *Mr1^+/+^* and *Mr1^-/-^*mice were stained for α-SMA (a) and Picrosirius Red (b), and representative images (10x) and quantification of positively stained areas are shown at the indicated time points. Fibrosis severity was independently evaluated across the entire liver using the Ishak scoring system by blinded board-certified pathologists. Fibrosis scores for individual animals are shown, alongside stacked bar graphs indicating the percentage of animals at each fibrosis score (0, 2, 2.5, 3, 3.5, 4) at days 12 and 26, based on a representative scoring assessment (c). To assess the contribution of MAIT cell activation, *Mr1^+/+^* mice were treated with the MR1 inhibitory ligand Ac-6-FP during CCl_4_ administration, and representative α-SMA staining (10x) and quantification are shown (d). All representative images are from the left hepatic lobe. Data are pooled from 4 (a-c) and 3 (d) independent experiments. Data are presented as mean ± SEM (a-d) and stacked bar plots (c). Statistical analyses were performed using two-way ANOVA for (a-c) and Mann–Whitney *U* tests for (d). * and ** denote differences with *p* < 0.05 and 0.01, respectively.

To test whether TCR-mediated MAIT cell activation underlies the enhanced fibrotic response in *Mr1*^+/+^ mice, we employed a pharmacological approach in which acetyl-6- formylpterin (Ac-6-FP), an MR1 inhibitory ligand^31^, was used to interfere with MAIT cell TCR signaling. Ac-6-FP administration attenuated α-SMA expression on day 12, effectively recapitulating the reduced fibrosis observed in *Mr1^-/-^* mice at this time point, but not on day 26 **(Fig. 2d)**. These findings implicate TCR-triggered MAIT cell activation as a key driver of early fibrotic remodeling.

Collectively, the above data demonstrate that MAIT cells amplify hepatocellular injury, myofibroblast activation, and matrix deposition during the early phase of chronic liver injury.

### Hepatic tissue injury alters MAIT cell residency and trafficking properties

We next sought to determine whether chronic liver injury changes the phenotypic characteristics of hepatic MAIT cells. Cytofluorimetric analyses **(Supplementary Fig. 3)** showed unchanged liver MAIT cell frequencies and absolute numbers when corn oil- and CCl_4_-treated mice were compared at acute (day 1), early (day 12), or established (day 26) stages of liver injury **(Fig. 3a, Supplementary Fig. 4a-b)**. Interestingly, however, the proportion of CD69^+^ MAIT cells was substantially reduced on both days 12 and 26, while the geometric mean fluorescence intensity (gMFI) of CD69 remained unchanged **(Fig. 3b)**. Notably, this reduction in CD69^+^ cells was evident as early as 24 hours following a single CCl_4_ injection **(Supplementary Fig. 4c)**. In contrast, CD25 expression was largely stable across all time points, with only a modest increase in CD25 gMFI on day 26 **(Fig. 3c, Supplementary Fig. 4d)**.

**Fig. 3.**
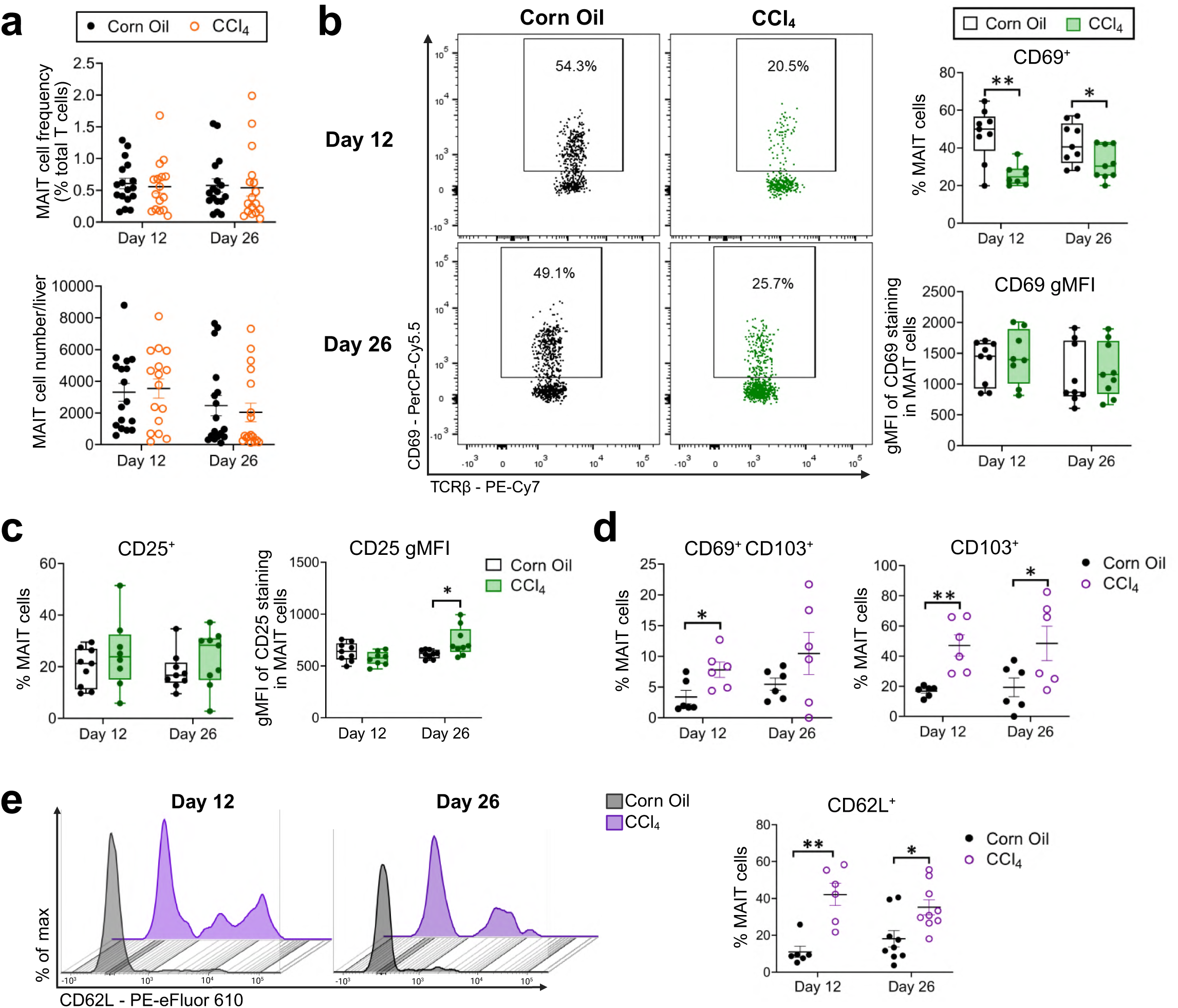
Tissue injury alters the residency-associated phenotype of hepatic MAIT cells without affecting their abundance. Corn oil- or CCl_4_-treated *Mr1^+/+^* mice were euthanized on days 12 or 26, and non-parenchymal HMNCs were isolated for flow cytometric analysis. The frequency of hepatic MAIT cells among T cells and absolute MAIT cell numbers were assessed at both time points (a). CD69 (b) and CD25 (c) expression on MAIT cells were examined by measuring the frequency of positive cells and the staining intensity (gMFI), with representative flow cytometry dot plots shown for CD69. Expression of residency-associated markers was also investigated by quantifying CD69^+^ CD103^+^ and CD103^+^ MAIT cell populations (d). CD62L^+^ MAIT cells were also examined, and representative overlaid histograms are depicted in (e). Data are pooled from 6 (a), 3 (b-c, e), and 2 (d) independent experiments. Data are presented as mean ± SEM (a, d-e) and box-and-whisker plots showing median, IQR, and min-max (b-c). Statistical analyses were performed using Student’s *t* tests or Mann–Whitney *U* tests. * and ** denote differences with *p* < 0.05 and 0.01, respectively.

The above changes were selective for MAIT cells because hepatic T_conv_ cells did not show a similar reduction in CD69. In fact, they exhibited a slight rise in CD69 by day 26 post-CCl_4_ **(Supplementary Fig. 5a)**. Of note, in our analyses, we defined T_conv_ cells as non-invariant natural killer T (*i*NKT), non-MAIT αβ TCR^+^ cells, thereby excluding *i*NKT and γδ T cell populations, which comprise a substantial proportion of T cells in the mouse liver^32^. T_conv_ cell abundance and CD25 expression also did not differ between treatment groups **(Supplementary Fig. 5b-c)**. Similarly, CD69 and CD25 expression on splenic MAIT cells was unaffected **(Supplementary Fig. 5d-e)**, indicating that these phenotypic changes were restricted to the hepatic compartment.

Since CD69 is closely linked to tissue retention^33,34^ and may not merely reflect T cell activation in chronic inflammatory settings, we next investigated whether chronic liver injury alters the tissue-residency characteristics of MAIT cells. Tissue-resident memory T_conv_ cells are commonly identified by virtue of their CD69 and CD103 co-expression. Despite an overall decline in CD69⁺ hepatic MAIT cells **(Fig. 3b)**, the proportion of CD103⁺CD69⁺ MAIT cells was increased in CCl_4_-treated mice on day 12, with a similar but non-significant trend on day 26 **(Fig. 3d)**. When CD103 was assessed independently, CCl_4_ treatment also led to a significant rise in CD103⁺ MAIT cells at both time points **(Fig. 3d)**. These changes were specific to MAIT cells as no pronounced differences in CD103⁺CD69⁺ or CD103⁺ T_conv_ cell frequencies were detectable **(Supplementary Fig. 5f-g)**. To further probe the migratory and residency potentials of MAIT cells, we assessed L-selectin (CD62L), an adhesion molecule involved in lymphocyte trafficking that may promote the progression of fibrosis^35^. We found CD62L-expressing MAIT cells to be more abundant in the liver on both days 12 and 26 following CCl_4_ injection **(Fig. 3e)**.

Taken together, the above findings indicate that hepatic MAIT cells undergo dynamic phenotypic changes, giving rise to distinct residency- and trafficking-associated features in the injured liver.

### Chronic liver injury gives rise to an exhaustion-associated phenotype without functionally impairing hepatic MAIT cells

In the absence of overt increases in activation markers, we set out to examine whether chronic liver injury induces chronic stimulation and exhaustion in MAIT cells. Hepatic MAIT cells from CCl_4_-treated mice exhibited a significant increase in the frequency of PD-1^+^ cells on both days 12 and 26, while PD-1’s gMFI was not affected **(Fig. 4a)**. This upregulation was not observed acutely at 24 hours **(Supplementary Fig. 6a)** and appeared to be restricted to the liver as splenic MAIT cells showed only a modest increase in PD-1 expression on days 12 and 26 that did not reach statistical significance **(Supplementary Fig. 6b)**. Also, importantly, this effect was MAIT cell-selective since T_conv_ cells showed no differences in PD-1 expression following CCl_4_ administration **(Fig. 4b)**.

**Fig. 4.**
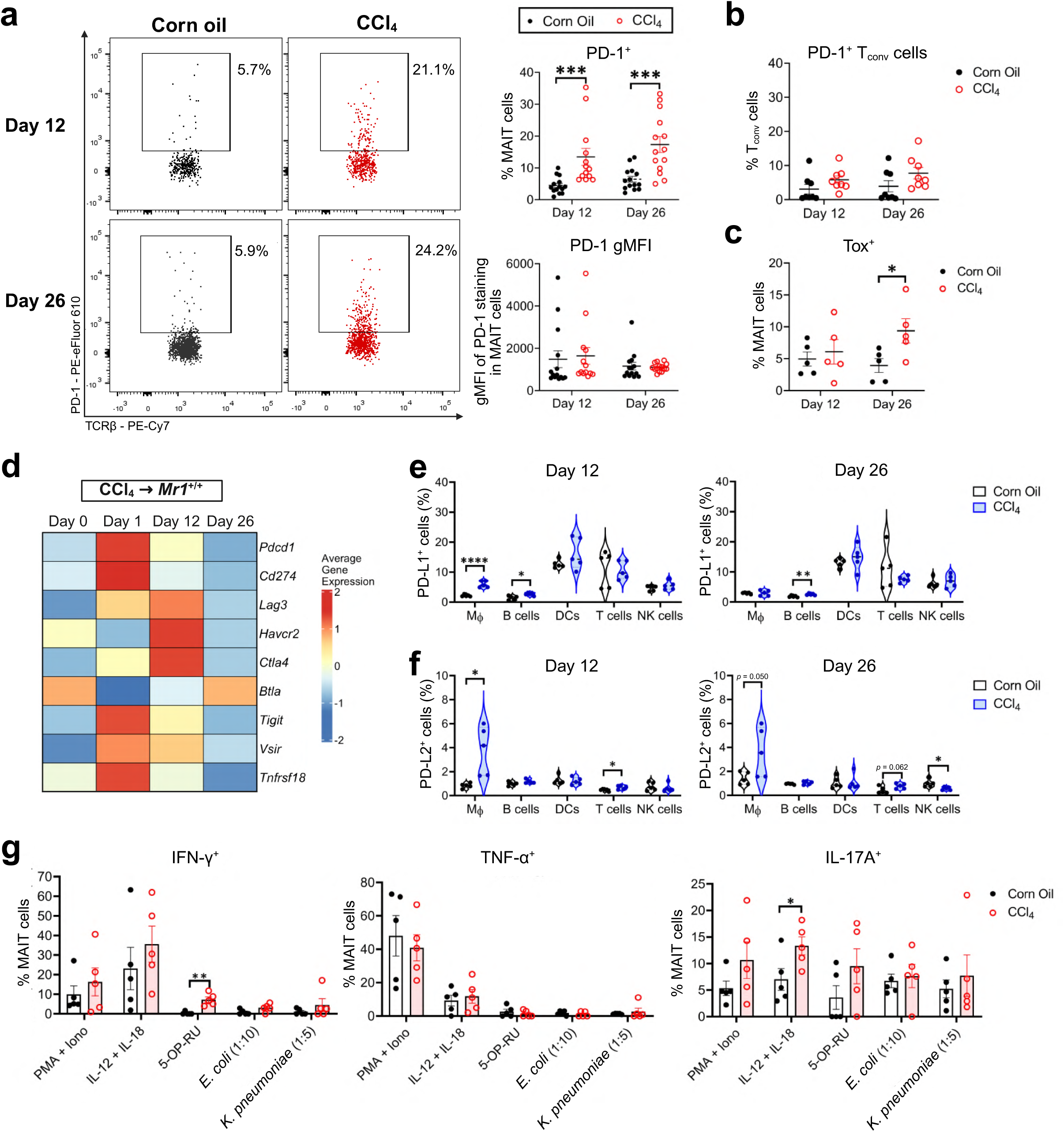
MAIT cells upregulate exhaustion-associated markers during early chronic liver injury while retaining effector function. PD-1 expression on hepatic MAIT cells was assessed by flow cytometry at days 12 and 26 and is presented as the frequency of PD-1^+^ cells and PD-1 gMFI, with representative dot plots (a). PD-1 expression on T_conv_ (5-OP-RU-loaded mouse MR1 tetramer^−^, PBS-57-loaded mouse CD1d tetramer^−^, TCRβ^+^) cells (b) and Tox expression on MAIT cells (c) were similarly evaluated at the indicated time points. Following bulk RNA-seq of HMNCs from CCl_4_-treated *Mr1^+/+^* mice, mean expression of exhaustion/dysfunction-associated genes was assessed and depicted as a heatmap (d). Expression of PD-L1 (e) and PD-L2 (f) across hepatic immune cell subsets (macrophages, B cells, DCs, T cells, and NK cells) was additionally measured flow cytometrically at days 12 and 26. IFN-γ, TNF, and IL-17A production by hepatic MAIT cells isolated at day 26 was interrogated by intracellular cytokine staining following *ex vivo* stimulation with PMA and ionomycin, IL-12 and IL-18, 5-OP-RU, *E. coli* lysate, or *K. pneumoniae* lysate (g). Data are pooled from 5 (a), 3 (b), and 2 (c, e-g) independent experiments. Data are presented as mean ± SEM (a-c, g) and violin plots showing distribution, median, and quartiles (e-f). Statistical analyses were performed using Student’s *t* tests or Mann-Whitney *U* tests. *, **, ***, and **** denote differences with *p* < 0.05, 0.01, 0.001, and 0.0001, respectively.

To extend our studies beyond PD-1 expression, we assessed Tox, a transcription factor mediating exhaustion and adaptation in chronically stimulated T_conv_ cells^36^. By day 26, MAIT cells from CCl_4_-treated mice exhibited elevated Tox levels **(Fig. 4c)**. Moreover, bulk RNA-seq analyses on HMNCs from *Mr1*^+/+^ mice revealed concomitant upregulation of multiple exhaustion-associated genes (*Pdcd1*, *Cd274*, *Lag3*, *Havcr2*, *Ctla4*, *Btla*, *Tigit*, *Vsir*, *Tnfrsf18*) across several time points after CCl_4_ treatment **(Fig. 4d)**, indicating the presence of a broad exhaustion-related signature within the injured liver compartment.

Because inhibitory receptor signaling depends on ligand availability, we examined the expression of PD-L1 and PD-L2. Both the frequency of PD-L1⁺ and PD-L2⁺ cells and the expression level of these molecules were increased in multiple hepatic immune subsets, including macrophages, B cells, T cells and NK cells, following CCl_4_ treatment **(Fig. 4e-f, Supplementary Fig. 6c-d)**. In contrast, PD-L1 and PD-L2 levels remained largely unchanged in splenic cell populations **(Supplementary Fig. 6e-f).** Notwithstanding, the protein expression of other inhibitory receptors, including LAG-3, TIM-3, CTLA-4, and TIGIT, was not markedly altered on hepatic MAIT cells post-CCl_4_ **(Supplementary Fig. 7a)**. Similarly, the expression of the death receptor Fas and the survival marker CD127 on MAIT cells was unaltered **(Supplementary Fig. 7b-c)**, suggesting that PD-1 upregulation was not associated with impaired survival or increased susceptibility to apoptosis.

Finally, to determine whether the observed exhaustion-associated phenotype translated into MAIT cell dysfunctions, we assayed for effector cytokine production in response to a panel of stimuli with different modes of action. To this end, HMNCs were incubated *ex vivo* with phorbol 12-myristate 13-acetate (PMA) and ionomycin to bypass TCR ligation leading to cellular activation, with IL-12 and IL-18 to drive cytokine-mediated activation, and with 5-OP-RU, *Escherichia coli* lysate or *Klebsiella pneumoniae* lysate to assess MR1-dependent MAIT cell responses. At the most advanced stage of fibrosis (day 26), MAIT cells from CCl_4_-treated mice produced IFN-γ, TNF and IL-17A at levels equal to or even greater than those detected in corn oil-injected controls in all of the above stimulation cultures **(Fig. 4g)**. A similar pattern was also evident at the earlier day-12 time point **(Supplementary Fig. 7d)**, indicating that MAIT cell effector function was maintained or even heightened over the course of liver injury. Therefore, although chronic liver injury promotes the acquisition of an exhaustion-associated phenotype in hepatic MAIT cells, the effector capacity of these cells remains intact, supporting their functional adaptation rather than terminal exhaustion.

### CCl_4_-elicited hepatic fibrosis is accompanied by robust MAIT17 polarization in the liver

MAIT cells in mice are typically divided into two functional subsets, namely MAIT1 and MAIT17 cells, based on their type-1 and type-17 cytokine production profiles, respectively. Given the association of type-17 immunity with fibrotic pathology^37–39^, we examined whether fibrosis alters hepatic MAIT cell polarization transcriptionally or functionally.

Chronic CCl_4_ exposure induced a strong shift towards a MAIT17 phenotype in the liver, characterized by a significant increase in MAIT17 (T-bet^−^RORγt^+^) cells and a corresponding decline in MAIT1 (T-bet^+^RORγt^−^) cells on day 12, with similar trends persisting on day 26 **(Fig. 5a)**. This MAIT1-to-MAIT17 shift was detectable as early as 24 hours following a single CCl_4_ injection **(Supplementary Fig. 8a)**. In contrast, spleens contained a less prominent MAIT17 cell population, while a shift towards a double-negative (T-bet^−^RORγt^−^) phenotype was manifest on day 12 **(Supplementary Fig. 8b).** Therefore, MAIT17 polarization appeared to be selective to the liver during CCl_4_ injury. Moreover, consistent with the observed decline in hepatic MAIT1 cells, the expression of CD122, a surrogate marker of MAIT1 cells^40,41^, was decreased, notably on day 26 and to a lesser extent on day 12 in the liver **(Fig. 5b)**.

**Fig. 5.**
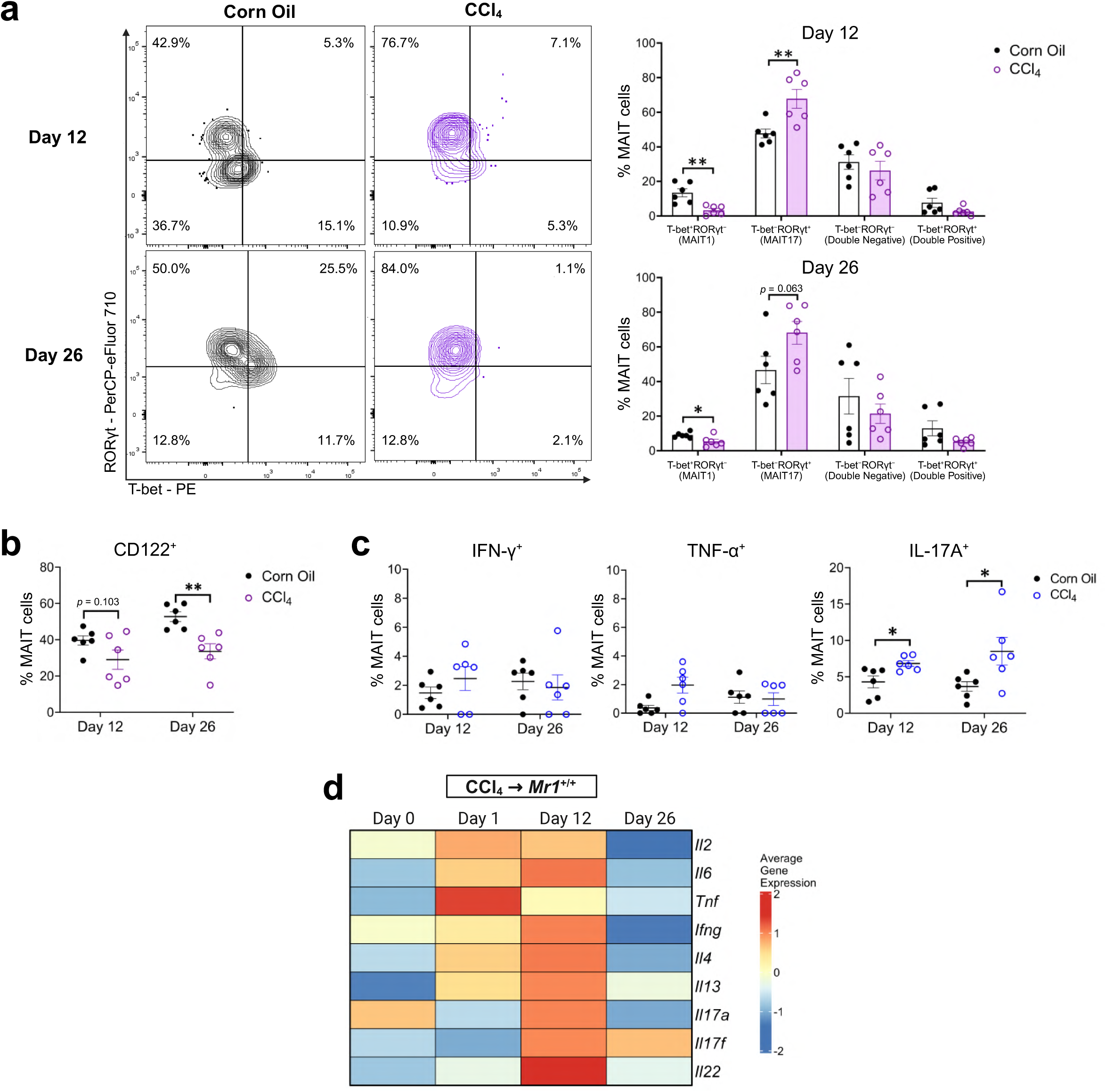
Hepatic MAIT cells skew toward a MAIT17 phenotype and exhibit increased IL-17A production during liver injury. Expression of T-bet and RORγt in hepatic MAIT cells was assessed by flow cytometry at the indicated time points and is presented as the frequency of T-bet^+^RORγt^−^ (MAIT1), T-bet^−^RORγt^+^ (MAIT17), T-bet^−^RORγt^−^ (Double Negative), and T- bet^+^RORγt^+^ (Double Positive) populations, with representative contour plots (a). CD122 expression on hepatic MAIT cells at the given time points was also quantified (b). Cytokine production by hepatic MAIT cells at days 12 and 26 in the absence of *ex vivo* stimulation is presented for IFN-γ, TNF, and IL-17A (c). Mean expression of cytokine genes in HMNCs from CCl_4_-treated *Mr1^+/+^* mice, derived from bulk RNA-seq data as described previously, is depicted (d). Data in (a-c) are pooled from 2 independent experiments. Data are presented as mean ± SEM. Statistical analyses were performed using Student’s *t* tests or Mann-Whitney *U* tests. * and ** denote differences with *p* < 0.05 and 0.01, respectively.

To functionally validate the observed phenotypic polarization, we assayed for MAIT cells’ cytokine content without any *ex vivo* stimulation. To this end, freshly isolated hepatic MAIT cells from CCl_4_-treated mice exhibited higher baseline IL-17A production on both days 12 and 26 compared to corn oil controls, reflecting a MAIT17-skewed profile **(Fig. 5c)**. In contrast, only negligible differences were detectable in IFN-γ^+^ or TNF^+^ cell proportions **(Fig. 5c),** and cytokine production by splenic MAIT cells remained unchanged **(Supplementary Fig. 8c)**. Complementing these findings, subjecting HMNCs to bulk transcriptomic profiling showed an enrichment of type-17 cytokine transcripts, including *Il17a*, *Il17f* and *Il22* as fibrosis progressed **(Fig. 5d).**

Taken together, these findings demonstrate that chronic liver injury reprograms hepatic MAIT cells toward a robust MAIT17 phenotype. Given the established role of IL-17-producing lymphocytes in fibrogenesis^37–39^, this shift represents a potential mechanism by which MAIT cells contribute to profibrotic remodeling in chronic liver injury.

### MAIT cells impede hepatic T_reg_ cell accumulation during chronic liver injury

MAIT cells are known to engage in crosstalk with multiple other cell types. Therefore, we investigated whether they shape the immune landscape of the injured liver by enumerating other immunocytes in *Mr1*^+/+^ and *Mr1*^-/-^ mice treated with CCl_4_ or corn oil. No substantial differences were found in the frequencies of B cells, NK cells, macrophages, dendritic cells (DCs), or myeloid-derived suppressor cells (MDSCs) between the two strains **(Supplementary Fig. 9a-e)**. Interestingly, however, a clear difference was notable in the T_reg_ cell compartment. Accordingly, *Mr1*^+/+^ mice displayed lower hepatic T_reg_ cell frequencies and absolute numbers on both days 12 and 26 after CCl_4_ treatment compared to Mr*1^-/-^*mice **(Fig. 6a-b)**. Consistent with these findings, bulk RNA-seq analyses revealed that T_reg_ cell-associated genes (*Foxp3*, *Il2ra*, *Ctla4*, *Ikzf2*, *Ikzf4*, *Il10*) were expressed at lower levels within CCl_4_-treated *Mr1*^+/+^ mouse livers compared with *Mr1*^-/-^ livers at all tested time points **(Fig. 6c)**. Moreover, when integrated into a composite transcriptional T_reg_ cell score, which we describe in Methods, this signature was weakened in *Mr1^+/+^* mice **(Fig. 6d)**. Again, the observed difference in T_reg_ cell abundance was liver-specific and not evident in the spleen **(Fig. 6e)**.

**Fig. 6.**
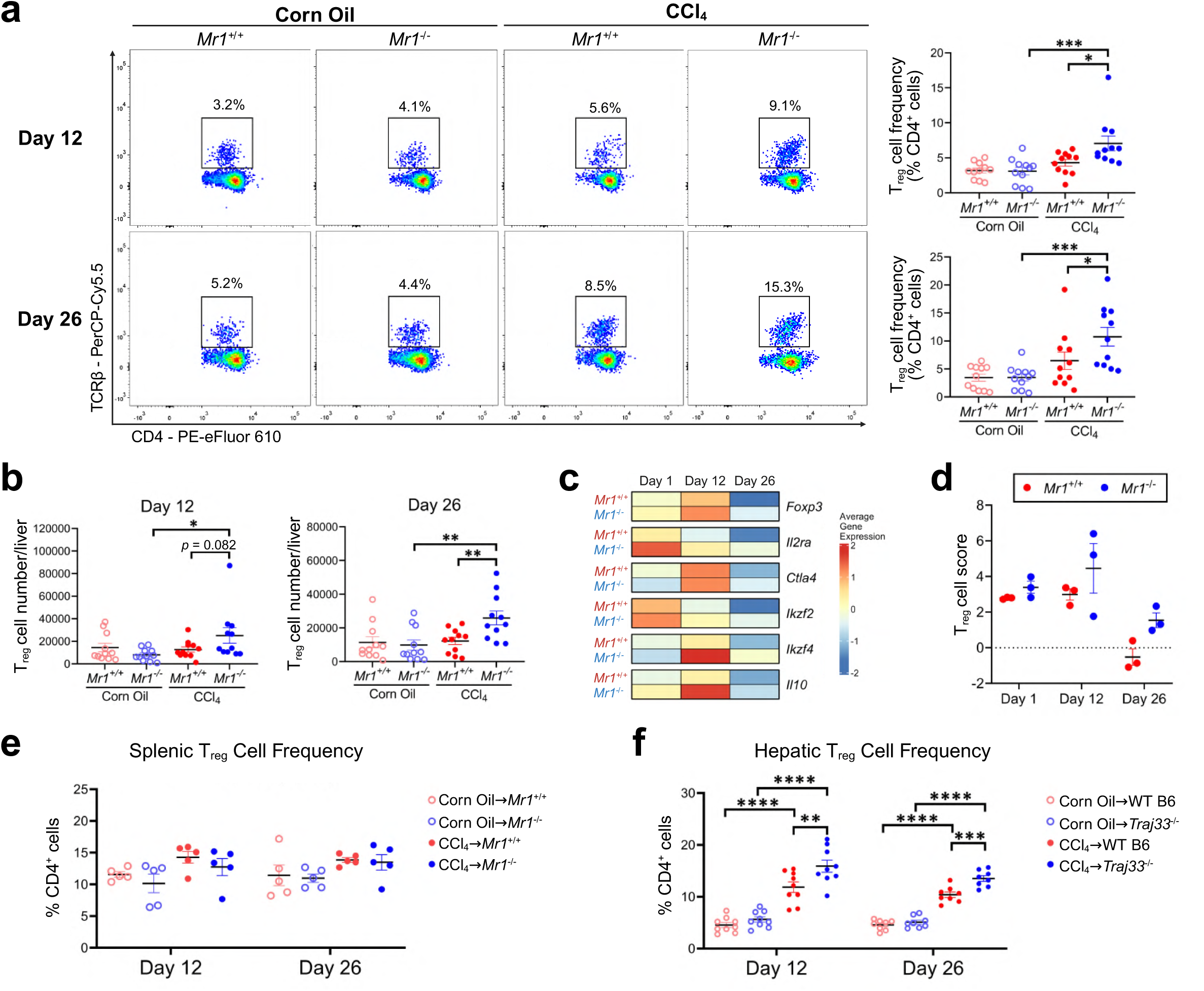
MAIT cells limit T_reg_ cell accumulation in the liver during injury. T_reg_ cell frequency among hepatic CD4*^+^*T cells was cytofluorometrically quantified in corn oil- or CCl_4_-treated *Mr1^+/+^*and *Mr1^-/-^* mice at days 12 and 26, with representative flow cytometry plots shown (a). Total hepatic T_reg_ cell numbers were also determined at the same time points (b). Mean expression of T_reg_-associated genes in non-parenchymal HMNCs from CCl_4_-treated *Mr1^+/+^* and *Mr1^-/-^* mice across days 1, 12, and 26 is depicted from bulk RNA-seq data (c), and composite T_reg_ cell scores calculated from these genes are shown across the same time points (d). T_reg_ cell frequency in the spleen was measured by flow cytometry in corn oil- or CCl_4_-treated *Mr1^+/+^* and *Mr1^-/-^* mice (e). Hepatic T_reg_ cell frequency was also assessed in MAIT cell-sufficient wildtype B6 and MAIT cell-deficient *Traj33^-/-^* mice following corn oil or CCl_4_ treatment at the indicated timepoints (f). Data are pooled from 4 (a-b) and 2 (e-f) independent experiments. Data are presented as mean ± SEM. Statistical analyses were performed using two-way ANOVAs. *, **, ***, and **** denote differences with *p* < 0.05, 0.01, 0.001, and 0.0001, respectively.

Our findings thus far demonstrated that MR1 deficiency leads to increased hepatic T_reg_ cell frequencies after CCl_4_ administration, which may serve to protect against liver injury, which is supported by less severe fibrosis in *Mr1^-/-^* mice **(Fig. 2a-c)**. This is also consistent with our observation that pharmacological inhibition of MAIT cell activation in *Mr1*^+/+^ animals attenuates liver fibrosis **(Fig. 2d)**, countering the remote possibility that MR1 deficiency fortuitously dictates the observed phenomena in a MAIT cell-independent fashion. Nonetheless, to further validate our results, we resorted to *Traj33*^-/-^ mice that are MR1-sufficient but devoid of MAIT cells due to impaired invariant TCR α-chain rearrangement^42^. Following CCl_4_ exposure, *Traj33*^-/-^mice exhibited significantly higher hepatic T_reg_ cell frequencies compared to wildtype B6 controls **(Fig. 6f)**, confirming that MAIT cell presence is associated with a downsized hepatic T_reg_ cell population during chronic liver injury.

We next asked whether MAIT cells impacted phenotypic and/or functional T_reg_ cell characteristics. Using high-dimensional flow cytometry, we examined the expression of eight T_reg_ cell-associated proteins (CD73, PD-1, GITR, CD39, HELIOS, CTLA-4, IL-10, LAP) on days 12 and 26 **(Fig. 7a-b)**. CCl_4_-treated mice exhibited increased CD39, CTLA-4, PD-1 and HELIOS levels on their T_reg_ cells compared to corn oil controls, indicative of an activated T_reg_ cell phenotype with increased suppressive potentials **(Fig. 7a-b and Supplementary Fig. 10a)**. However, no major differences were detected in these markers between *Mr1*^+/+^ and *Mr1*^-/-^ mice. Therefore, MAIT cells simply alter hepatic T_reg_ cell numbers, not their immunosuppressive behaviors at an individual cell level.

**Fig. 7.**
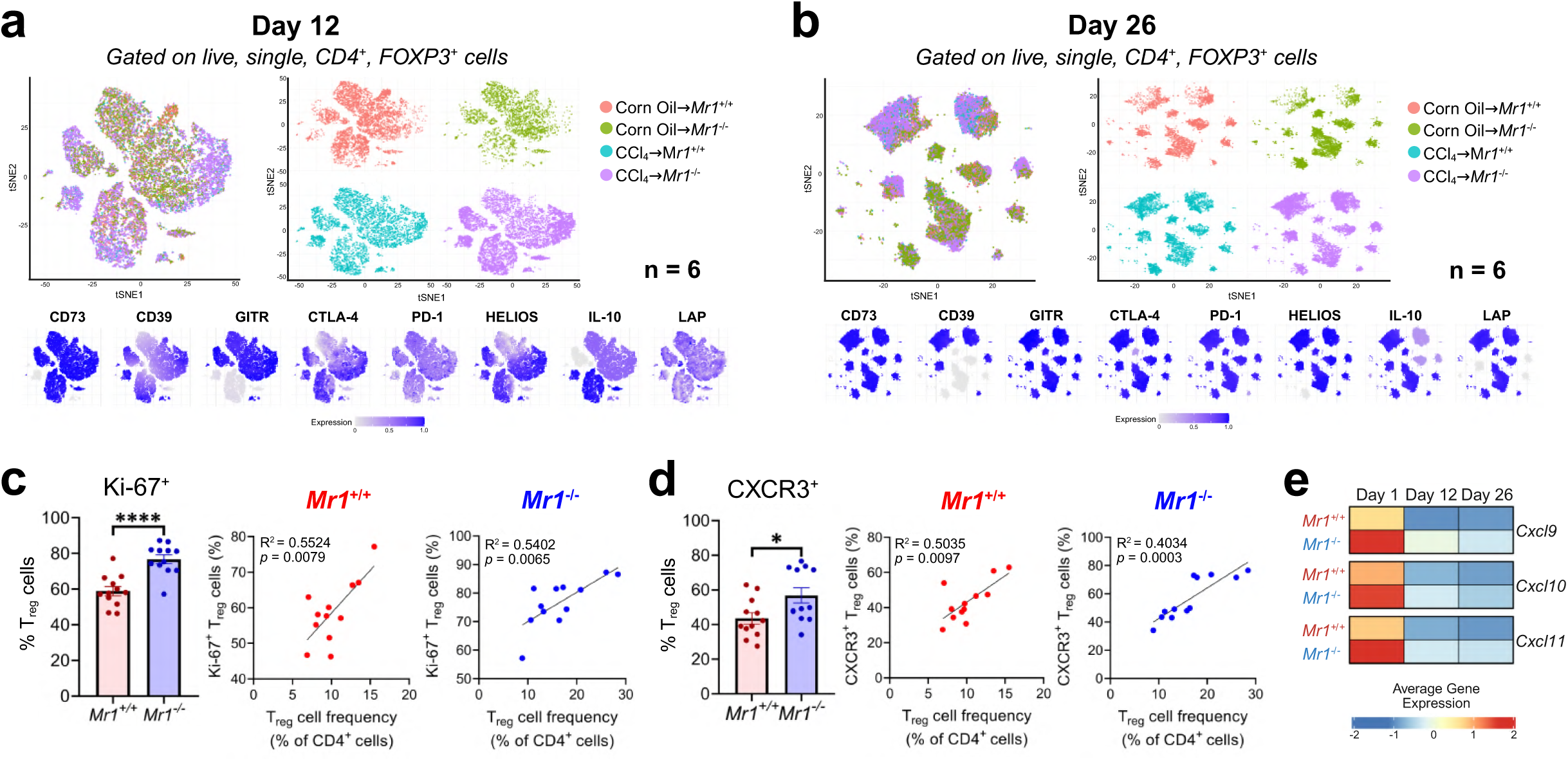
Hepatic T_reg_ cells in *Mr1*^-/-^ mice exhibit enhanced proliferation and chemokine receptor expression during chronic injury. High-parameter flow cytometry was performed on hepatic T_reg_ cells (live, single, CD4^+^, FoxP3^+^) from corn oil- or CCl_4_-treated *Mr1^+/+^* and *Mr1^-/-^*mice at days 12 (a) and 26 (b) and visualized by t-SNE with expression of CD73, CD39, GITR, CTLA-4, PD-1, HELIOS, IL-10, and LAP projected onto the shared embedding. Ki-67 (c) and CXCR3 (d) expression at day 12 are presented as the frequency of Ki-67^+^ and CXCR3^+^ T_reg_ cells, respectively, along with their relationships to total hepatic T_reg_ cell frequency in *Mr1^+/+^* and *Mr1^-/-^* mice. Mean expression of *Cxcl9*, *Cxcl10*, and *Cxcl11* in non-parenchymal HMNCs from CCl_4_-treated *Mr1^+/+^* and *Mr1^-/-^* mice is depicted from bulk RNA-seq data (e). Data are pooled from 2 independent experiments. Data are presented as mean ± SEM. Statistical analyses were performed using a Student’s *t* test (c, left), a Mann-Whitney *U* test (d, left), and linear regression (c,d, right). * and **** denote differences with *p* < 0.05 and 0.0001, respectively.

To explore the mechanisms underlying strain differences in T_reg_ cell frequencies, we analyzed T_reg_ cell proliferation, trafficking and apoptosis. These experiments demonstrated higher Ki-67^+^ and CXCR3^+^ T_reg_ cell frequencies in CCl_4_-treated *Mr1*^-/-^ mice, both of which also positively correlated with overall hepatic T_reg_ cell abundance **(Fig. 7c-d)**. Similar increases were observed within the total CD4^+^ T cell compartment **(Supplementary Fig. 10b)**. In line with this observation, our bulk RNA-seq analyses demonstrated higher transcript levels for CXCR3- associated chemokines *Cxcl9*, *Cxcl10* and *Cxcl11* in *Mr1*^-/-^ livers on days 1, 12 and 26 **(Fig. 7e)**, with *Cxcl10* being significantly elevated at early time points (days 1 and 12) **(Fig. 1e)**. Therefore, increased T_reg_ cell proliferation and their concomitant CXCR3-mediated recruitment appear to contribute to their intrahepatic accumulation in the absence of MAIT cells. In contrast, no difference in overall apoptosis was detectable, as judged by intracellular cleaved caspase-3 levels **(Supplementary Fig. 10c)**. Interestingly, T_reg_ cells from *Mr1*^-/-^ mice displayed reduced levels of the anti-apoptotic factor Bcl-2 and increased expression of the pro-apoptotic receptor Fas (CD95) compared to *Mr1*^+/+^ animals **(Supplementary Fig. 10d)**.

To confirm that the proposed MAIT cell-T_reg_ cell axis contributes to fibrogenesis, we injected mice with PC61 **(Fig. 8a)**, an anti-CD25 monoclonal antibody known to inactivate T_reg_ cells^43^. PC61 pre-treatment resulted in a near-complete removal of CD4^+^CD25^+^FoxP3^+^ cells in both *Mr1*^+/+^ and *Mr1*^-/-^ mice **(Fig. 8b)**. As expected, isotype-treated *Mr1*^+/+^ and *Mr1*^-/-^ mice recapitulated strain-dependent differences in hepatic T_reg_ cell frequencies **(Fig. 6a and 8b)**. In the next series of experiments, we examined the severity of hepatic fibrosis via α-SMA staining in animals with or without T_reg_ cell inactivation. Among isotype-treated animals, *Mr1*^+/+^ mice again exhibited greater α-SMA⁺ areas compared to *Mr1*^-/-^ mice on day 12, consistent with our earlier findings **(Fig. 2c and 8c)**. However, and as hypothesized, the difference in fibrosis was eliminated following PC61 treatment, indicating that T_reg_ cells were required for MAIT cell-accelerated fibrosis **(Fig. 8c)**.

**Fig. 8.**
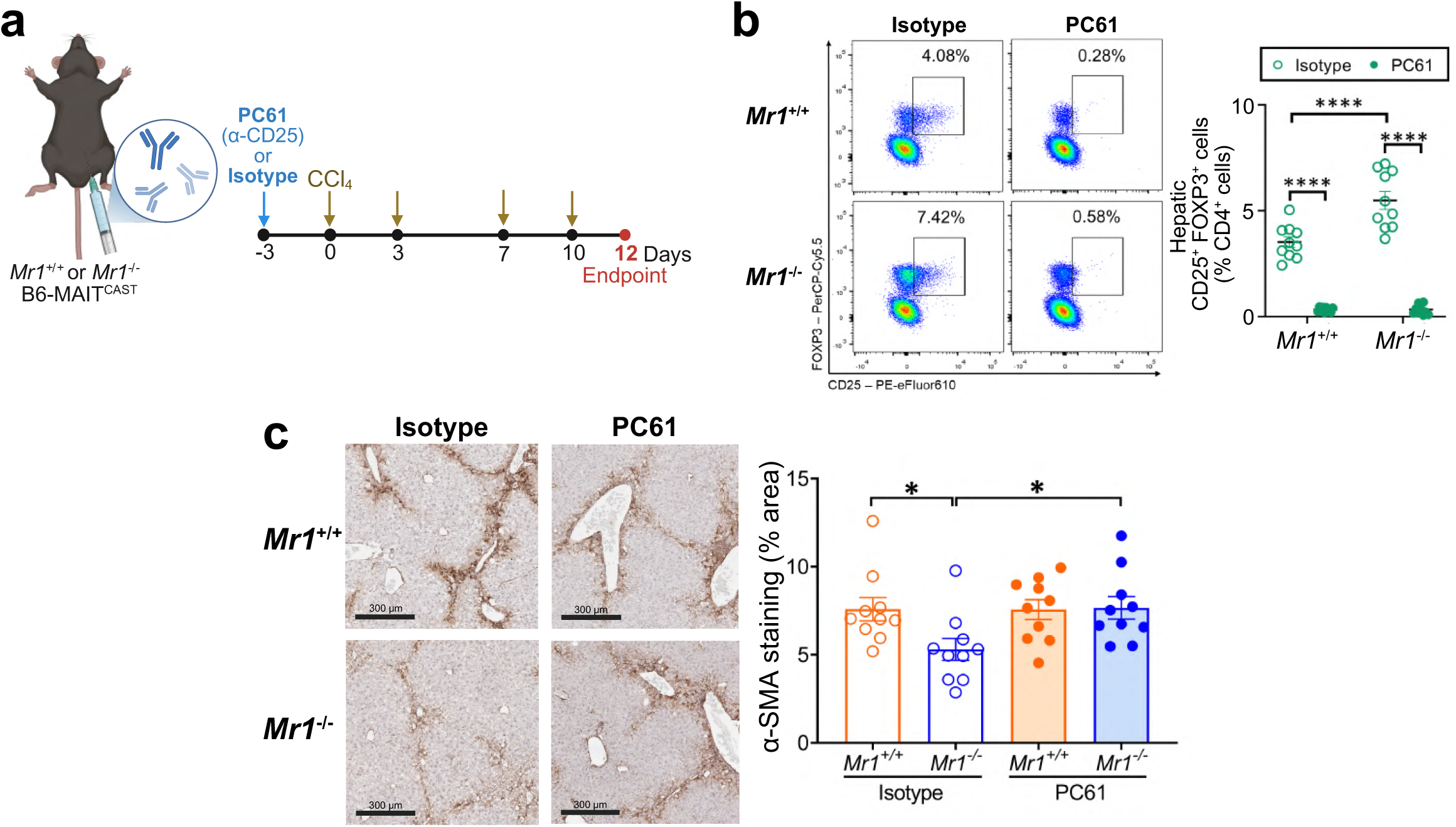
T_reg_ cell inactivation abrogates MAIT cell-driven pro-fibrotic effects. An anti-CD25 monoclonal antibody (PC61) or isotype control was administered to *Mr1^+/+^* or *Mr1^-/-^* mice prior to CCl_4_ treatment to inactivate T_reg_ cells, and mice were analyzed at day 12 (a). Hepatic CD4^+^ CD25^+^ FoxP3^+^ T_reg_ cell frequency was quantified by flow cytometry following treatment, with representative flow cytometry plots shown in (b). Liver fibrosis was assessed by α-SMA staining in CCl_4_-treated *Mr1^+/+^* or *Mr1^-/-^* mice and is presented as the proportion of α-SMA⁺ area (c). Data are pooled from 2 independent experiments. Data are presented as mean ± SEM. Statistical analyses were performed using two-way ANOVAs. * and **** denote differences with *p* < 0.05 and 0.0001, respectively.

### Human MAIT cells from fibrotic livers exhibit pro-fibrotic, MAIT17-biased and exhaustion-associated transcriptomic signatures

To determine whether the phenotypic changes observed in our mouse models hold true in human liver fibrosis, we analyzed a publicly available single-cell RNA-seq dataset of human liver CD45^+^ cells^44^ **(Fig. 9a)**. From this dataset, we included all samples with available fibrosis annotation, comprising five non-fibrotic and seven fibrotic liver samples spanning a range of severities, from mild pericellular fibrosis to cirrhosis **(Supplementary Table 1)**. We found that a cell cluster originally annotated as circulating effector memory T (T_EM_) cells by Guilliams *et al.*^44^ was highly enriched for canonical MAIT cell markers, including *SLC4A10*, *ZBTB16*, *KLRB1*, *NCR3*, *IL23R*, *ME1*, *LST1* and *TMIGD2* **(Fig. 9a; Supplementary Fig. 11a-c)**. This transcriptional profile closely matched our previously described MAIT cell gene signature^45^. This was not surprising because since MAIT cells are known to display an effector memory T cell-like phenotype^46^. The noted features strongly suggested that the population in question represented MAIT cells, which we analyzed as such in our downstream analyses. MAIT cells were detected in both fibrotic and non-fibrotic liver samples with similar abundance among total T cells **(Fig. 9b-c)**. This observation aligns with our mouse data, in which fibrotic injury was associated with phenotypic alterations in MAIT cells without changing their overall frequency **(Fig. 3a).**

**Fig. 9.**
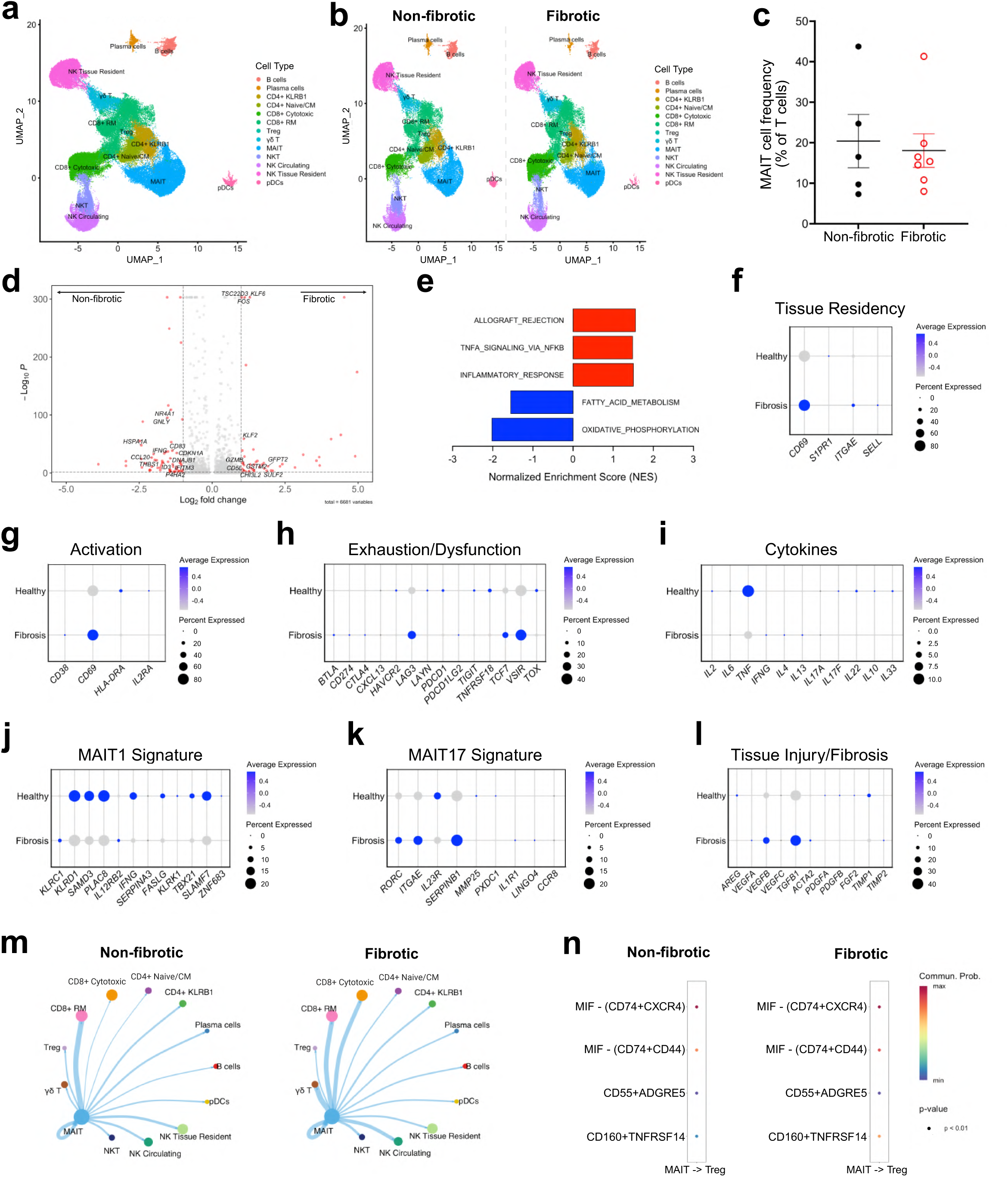
Human liver single-cell transcriptomics explores MAIT cell phenotypes in fibrotic and non-fibrotic conditions. Single-cell RNA-seq data from human liver CD45^+^ immune cells generated by Guilliams *et al.*^44^ were visualized using UMAP and annotated according to previously defined clusters (a). The circulating effector memory T (T_EM_) cell cluster was re- annotated as MAIT cells, and distribution across non-fibrotic and fibrotic samples is shown (b). MAIT cell abundance among T cells was quantified in non-fibrotic and fibrotic conditions (c). Differential gene expression between MAIT cells from fibrotic and non-fibrotic livers is presented as a volcano plot (d), and gene set enrichment analysis of these genes identified significantly enriched Hallmark pathways (e). MAIT cell transcriptional programs were further characterized using gene signature analysis, including tissue residency (f), activation (g), exhaustion/dysfunction (h), cytokine production (i), MAIT1-associated (j), MAIT17-associated (k), and tissue injury/fibrosis-related signatures (l). Inferred cell-cell communication networks between MAIT cells and other immune populations were analyzed using CellChat^47^, with interaction strength reflected by line thickness in non-fibrotic and fibrotic conditions (m). Predicted ligand-receptor interactions between MAIT and Treg cells are also shown (n).

Differential gene expression analyses revealed transcriptional changes in MAIT cells from fibrotic livers, including upregulated pro-inflammatory genes such as *Fos* and *Klf6* **(Fig. 9d)**. GSEA further demonstrated upregulated gene sets linked to inflammatory responses and TNF signaling via NF-κB in MAIT cells from fibrotic livers, along with reduced metabolic programs including fatty acid metabolism and oxidative phosphorylation **(Fig. 9e)**.

Consistent with our mouse findings, MAIT cells from fibrotic human livers exhibited altered gene expression associated with tissue residency and retention, including *CD69* and *ITGAE* (CD103), when compared to MAIT cells from non-fibrotic controls **(Fig. 9f)**. MAIT cells from fibrotic livers also showed reduced *HLA-DRA*, which is viewed as an activation marker **(Fig. 9g),** along with increased transcripts encoding multiple exhaustion- and dysfunction-associated genes, including *LAG3*, *VSIR* (VISTA), and *TCF7* **(Fig. 9h)**. The baseline expression of indicated cytokine genes was too low to allow for meaningful comparisons between non-fibrotic and fibrotic samples **(Fig. 9i)**. We also found MAIT cells to contain more *CD69* and less *LAG3* and *VSIR* transcripts when they were directly compared with CD8^+^ T_conv_ cells from fibrotic livers **(Supplementary Fig. 11d-f)**. Injury- and fibrosis-associated transcripts also differed between the two populations, with MAIT cells exhibiting higher *TNF* and *VEGFB*, while CD8⁺ T_conv_ cells showed elevated *TGFB1* **(Supplementary Fig. 11g-h)**.

Using this dataset, we also investigated whether MAIT cells exhibit MAIT17 polarization in liver fibrosis. As theorized, MAIT cells displayed reduced type-1 (e.g., *IFNG, TBX21*) and enriched type-17-associated gene transcripts, including elevated *RORC*, *ITGAE and SERPINB1* **(Fig. 9j–k)**. In addition, MAIT cells in fibrotic livers showed increased gene expression linked to tissue injury and fibrotic remodeling, including *VEGFB* and *TGFB1* **(Fig. 9l)**.

Finally, to explore potential interactions between MAIT cells and other immune cell populations, particularly T_reg_ cells, we performed ligand-receptor inference analysis using CellChat^47^. The T_reg_ cell cluster was confirmed by the expression of canonical T_reg_ markers, namely *CD4*, *FOXP3*, *CTLA4* and *IKZF2* **(Supplementary Fig. 11i)**. These analyses indeed revealed interactions between MAIT and T_reg_ cells in both non-fibrotic and fibrotic livers, primarily through pathways involving CD74-CXCR4 and CD74-CD44 axes **(Fig. 9m–n)** although these pathways were similarly represented in fibrotic and non-fibrotic conditions **(Fig. 9n)**.

Collectively, MAIT cells in fibrotic human livers acquire transcriptional features associated with chronic activation/dysfunction, MAIT17 polarization, and pro-fibrotic remodeling. Importantly, these signatures closely mirror those observed in our mouse models, supporting the translational relevance of our findings.

## DISCUSSION

MAIT cells are highly enriched in the liver and are important regulators of hepatic immunity; yet, their roles in various stages of liver fibrosis have been ill-defined. In this work, we identify MAIT cells as active contributors to early fibrotic remodeling, expanding their activities beyond those observed in advanced disease. Using a relatively comprehensive repertoire of MAIT cell-sufficient and -deficient models, including *Mr1*^+/+^, *Mr1*^-/-^ and *Traj33^-^*^/-^ mice, we demonstrate that MAIT cells promote fibrogenesis and are associated with reduced hepatic T_reg_ cell accumulation. These findings define a previously unrecognized MAIT cell-T_reg_ cell axis that can dramatically shape the injured liver milieu.

In contrast with studies of human fibrotic disease, in which MAIT cell depletion is frequently reported^14,15^, we found that MAIT cell abundance remained stable throughout fibrosis development. This discrepancy likely reflects differences in disease stage, as human studies are often restricted to biopsy or explanted tissue from patients with advanced disease.

Consistent with their innate-like capacity for rapid activation^48^, we found MAIT cells to exert their greatest influence during fibrosis initiation. Although we demonstrated that their association with fibrosis severity diminished at later stages, MAIT cells remained phenotypically and functionally active, as evidenced by sustained cytokine production and increased PD-1 expression in the absence of additional inhibitory receptors. PD-1 upregulation without a loss of function has similarly been reported for human intrahepatic MAIT cells in the context of cirrhosis^14^. These findings suggest that MAIT cells remain active during the later stages of fibrosis, but their relative contribution is diminished as additional immune and stromal pathways become dominant drivers of disease progression.

Mechanistically, our data support a role for MR1-dependent activation in driving MAIT cell-mediated fibrosis. We found that pharmacological inhibition of MR1 signaling with Ac-6-FP^31^ attenuated early fibrotic remodeling. This pathway is particularly relevant given the liver’s unique anatomical position within the gut-liver axis. Indeed, hepatic MAIT cells are continuously exposed to gut-derived microbial metabolites delivered via the portal vein, including riboflavin-derived ligands produced by bacteria such as members of the Bacilli and Proteobacteria classes^49,50^. These ligands can be directly presented by liver-resident cells, including hepatocytes and Kupffer cells, enabling local MR1-dependent activation^51,52^. During chronic liver injury, gut barrier dysfunction and microbial translocation further increase the availability of these ligands^53,54^, thereby amplifying MR1-dependent MAIT cell activation and promoting fibrogenic responses.

In parallel, chronic injury induced pronounced phenotypic changes within the hepatic MAIT cell compartment. We observed a rapid reduction in CD69 expression on MAIT cells, detectable as early as 24 hours post-injury. Commonly viewed as an early activation marker, CD69 also enables tissue retention^33,34^. We found the proportion of CD69⁺CD103⁺ MAIT cells to increase during fibrosis development, indicating the enrichment of a tissue-resident memory-like MAIT cell subset. Expansion of CD69⁺CD103⁺ MAIT cells has also been described in patients with primary biliary cholangitis, even as total intrahepatic MAIT cell frequencies decline^55^. Similar phenotypic changes have been reported for CD8⁺ T_conv_ cells in human liver disease, including reduced CD69 on circulating CD8⁺ T_conv_ cells in chronic hepatitis B^56^ and enrichment of CD69⁺CD103⁺ tissue-resident CD8⁺ T_conv_ cells in biliary atresia and primary biliary cholangitis^57,58^. In our model, however, changes in CD69 and CD103 were largely restricted to MAIT cells and were not mirrored by T_conv_ cells, underscoring the heightened sensitivity of MAIT cells to local cues within the injured liver.

CD69⁺CD103⁺ tissue-resident MAIT cell subsets are reportedly enriched for type-17 transcriptional programs in both steady-state and inflammatory settings^55,59,60^. In our model, pronounced polarization toward a MAIT17 phenotype was observed in the course of fibrosis development. Likewise, enrichment of IL-17^+^ MAIT cells has been reported in cirrhotic human livers^14^. We extend these findings by demonstrating that hepatic MAIT17 polarization emerges early during injury and is detectable even during acute liver injury, resembling other acute injury contexts such as skin wound healing^13^. Given that MAIT cells are recognized as important intrahepatic producers of IL-17A alongside Th17 cells^61^, this shift likely represents a significant source of type-17 inflammatory signaling in the liver. IL-17A can promote inflammatory responses and HSC activation, thereby enhancing collagen deposition^37–39^. Accordingly, disruption of IL-17 signaling has been shown to attenuate inflammation and fibrosis in multiple mouse models of liver injury^62–64^. Together, these findings suggest that MAIT17 polarization contributes to fibrotic remodeling in the injured liver.

Another central finding of this study is the identification of a MAIT cell-T_reg_ cell axis that regulates the severity of fibrosis. T_reg_ cells are well established as protective regulators of inflammation and fibrosis in toxin-induced liver injury models^62,65,66^. In the CCl_4_ model in particular, repeated injury has been shown to promote the expansion and recruitment of intrahepatic T_reg_ cells, which suppress inflammation and limit tissue damage and fibrogenesis^62,65,66^. While previous studies have shown that MAIT cells can interact with fibroblasts^14,15^, macrophages^16^, and NK cells^17^ in fibrotic settings, no study has directly examined how MAIT cells influence T_reg_ cells during fibrosis development despite prior discussion of potential MAIT-T_reg_ crosstalk in gut-liver inflammatory disease^67^.

Here, we provide experimental evidence that MAIT cell presence is associated with reduced hepatic T_reg_ cell accumulation during chronic injury. This was accompanied by increased frequencies of proliferating (Ki-67^+^) and CXCR3^+^ T_reg_ cells in MAIT cell-deficient *Mr1^-/-^*mice. In parallel, hepatic expression of CXCR3-associated chemokines (*Cxcl9*, *Cxcl10*, and *Cxcl11*) was elevated, supporting enhanced recruitment of T_reg_ cells in the absence of MAIT cells. Furthermore, although overall apoptosis was comparable at the population level, T_reg_ cells in *Mr1^-/-^* mice exhibited reduced Bcl-2 and increased Fas expression compared to *Mr1*^+/+^ mice.

These findings suggest that T_reg_ cells in *Mr1^-/-^* mice may exhibit increased susceptibility to cell death, potentially reflecting a compensatory mechanism that offsets their higher proliferative and recruitment capacity. Functionally, we also found T_reg_ inactivation to abrogate MAIT cell-dependent differences in fibrosis severity, establishing T_reg_ cells as critical downstream mediators of MAIT cell-driven fibrogenesis. To date, experimental evidence for MAIT cell-T_reg_ cell crosstalk in other settings remains limited. Previous work, including our own, has demonstrated that MAIT cells engage in crosstalk with T_reg_ cells in cancer settings. In bladder cancer, MAIT cell presence was associated with increased T_reg_ cell accumulation within the tumor microenvironment and exacerbated disease^19^. Similarly, MAIT cells have been shown to promote T_reg_ cell activity in hepatocellular carcinoma through TNF-TNFRSF1B signaling^68^. In contrast, our fibrosis model demonstrates the opposite relationship, with MAIT cells limiting T_reg_ cell accumulation. Together, these findings indicate that MAIT cells differentially regulate T_reg_ cell dynamics depending on disease context, promoting T_reg_ cell expansion in cancer while limiting T_reg_ cell accumulation during fibrotic injury, with detrimental consequences in both settings.

Lastly, to assess the translational relevance of our findings, we analyzed a publicly available human liver single-cell RNA-seq dataset^44^ with fibrosis scoring that did not rely on end-stage explant tissue from transplant patients. This allowed us to examine MAIT cells in a milder fibrotic environment that is more analogous to our experimental model. In this setting, human MAIT cells exhibited a transcriptional program that largely paralleled our mouse findings. One notable difference was a modest increase in CD69 expression in human MAIT cells from fibrotic samples compared with non-fibrotic counterparts, in contrast to our mouse model, where CD69 expression was reduced, potentially reflecting species-specific differences that warrant further investigation. Moreover, ligand-receptor inference analyses further predicted potential communication between MAIT and T_reg_ cells through CD74-CXCR4 and CD74-CD44 interactions. Notably, these predicted interactions were observed in both non-fibrotic and fibrotic liver samples, suggesting that these signaling pathways are not uniquely induced by fibrosis. These findings raise the possibility that MAIT cells may influence T_reg_ cell dynamics indirectly through interactions with other immune cell populations within the hepatic microenvironment, although this remains to be explored.

In summary, our study identifies MAIT cells as early contributors to liver fibrogenesis and links their activity to MR1-dependent activation, MAIT17 polarization, phenotypic adaptation, and modulation of hepatic T_reg_ cell responses. Therefore, MAIT cells can be viewed as an attractive target for anti-fibrotic therapies, and MAIT cell-directed interventions may be most effective when applied early in disease, during fibrotic initiation before fibrosis becomes established.

## METHODS

### Animals

All experiments were conducted using age- and sex-matched mice between 8 and 14 weeks of age. *Mr1*^+/+^ and *Mr1*^-/-^ B6-MAIT^CAST^ mice were kindly provided by Dr. Olivier Lantz (Curie Institute, Paris, France)^18^ and bred in house. *Traj33*^-/-^ mice have been previously described^42^. B6 mice were purchased from The Jackson Laboratory (strain #000664) and used as strain-matched wildtype controls for *Traj33*^-/-^ mice. All animals were bred and maintained under specific pathogen-free conditions at Western University’s West Valley Barrier and housed under a 12-hour light/dark cycle with *ad libitum* access to food and water. All experimental procedures were approved by the Western University Animal Care Committee under Animal Use Protocols 2022-0206 and 2024-0045.

### Mouse models of liver injury and fibrosis

To model liver injury and fibrosis, repeated liver damage was induced by *i.p.* injection of 0.6 µL CCl_4_ (Sigma-Aldrich) per gram of body weight diluted 10% v/v in corn oil (Sigma-Aldrich), administered twice weekly^16,22^. Control mice received an equivalent volume of corn oil. Mice were euthanized on Day 12 or Day 26, corresponding to the early and established phases of fibrosis, respectively, 48 hours after the final injection^21,22^. In the acute liver injury setting, mice received a single *i.p.* injection of CCl_4_ (0.6 µL/g body weight; 10% v/v in corn oil) and were euthanized 24 hours later.

### In vivo MAIT cell and T_reg_ cell inactivation protocols

To inactivate MAIT cells during chronic liver injury, animals were given 50 nmol Ac-6-FP (Cayman Chemicals) *i.p.* in 200 µL PBS on days -3, -1, and 0 relative to the start of the CCl_4_ regimen, and every other day thereafter^16,19,69^. Control animals received 200 µL PBS instead on the same schedule.

For T_reg_ cell inactivation, additional cohorts were injected *i.p.* with 1 mg anti-CD25 monoclonal antibody (clone PC61, Bio X Cell) 3 days before the initiation of CCl_4_ administration and euthanized on day 12 for endpoint analyses^70^. Control animals received 1 mg of a rat IgG1 isotype control (clone HRPN, Bio X Cell).

### Serum ALT quantification

Blood was collected from *Mr1*^+/+^ and *Mr1*^-/-^ B6-MAIT^CAST^ mice at endpoint by cardiac puncture. Serum was isolated by centrifugation (2000 × *g* for 15 minutes at 4°C) after clotting and analyzed for ALT levels using a mouse ALT ELISA kit (Abcam) according to the manufacturer’s instructions. Absorbance at 450 nm was measured using an Agilent BioTek Cytation 5 plate reader.

### Histopathology and image analysis

At indicated endpoints, livers were collected and excised into four major lobes (left, right, caudate and median). Tissues were fixed in 10% neutral-buffered formalin, processed, and embedded in paraffin wax. Liver sections were cut at 5-µm thickness and stained with hematoxylin and eosin (H&E), Masson’s Trichrome, and Picrosirius Red (Polysciences Inc.). Immunohistochemical staining for α-SMA was performed using a rabbit anti-α-SMA antibody (Abcam).

Stained slides were subjected to whole slide imaging using a Leica Aperio AT2 Brightfield Slide Scanner at 20× magnification and viewed with ImageScope (v12.4.6.5003). For each liver, five representative, non-overlapping fields were selected for quantitative image analysis. Unless otherwise indicated, all quantifications were performed on sections from the left hepatic lobe to maintain consistency across samples. In supplementary analyses, sections from individual liver lobes were quantified separately to assess potential regional differences. Tissue sections stained with Picrosirius Red and α-SMA were analyzed in ImageJ (v1.8.0_345) using the Colour Deconvolution 2 plugin to determine the percentage of positively stained areas. Thresholding parameters were standardized across all images within each staining type to ensure consistent quantification.

In addition to quantitative image analyses, tissue slides were independently evaluated by board-certified pathologists who were blinded to the experimental groups. Fibrosis severity was assessed across the entire liver using the Ishak fibrosis scoring system, a 7-tier histopathological score (0-6), where higher scores indicate more severe scarring^71^ based on Masson’s Trichrome-, H&E-, and Picrosirius Red-stained sections. Ballooning and inflammation were evaluated separately on H&E-stained sections and standard histopathological criteria.

### Tissue processing, single cell preparation and flow cytometric analysis

Livers were rinsed in cold PBS, minced, and homogenized to generate single-cell suspensions. The resulting cell preparations were washed twice with cold PBS and subjected to density gradient centrifugation using 33.75% Percoll PLUS (Cytiva) at 700 × *g* for 12 minutes at room temperature with no brake to enrich for mononuclear cells. Pelleted cells were then treated with ACK lysis buffer to remove residual erythrocytes. Spleens were similarly homogenized, centrifuged, and subjected to ACK lysis to obtain single-cell suspensions.

Isolated cells were then incubated with 5 µg/mL of an Fcγ receptor-blocking monoclonal antibody (anti-mouse CD16/CD32, clone 2.4G2) for 10 minutes at 4°C to prevent non-specific antibody binding. Surface staining was performed at 4°C for 30 minutes, except for samples stained with tetramers, which were incubated at room temperature to optimize tetramer binding. A complete list of monoclonal antibodies used for staining is provided in **Supplementary Table 2**. 5-OP-RU-loaded MR1 tetramer, 6-FP-loaded MR1 tetramer (staining control), PBS57-loaded CD1d tetramer, and unloaded CD1d tetramer (staining control) reagents were provided by the NIH Tetramer Core Facility (NIH Contract 75N93020D00005 and RRID:SCR_026557; Atlanta, USA).

For intracellular staining, cells were fixed and permeabilized using the FoxP3/Transcription Factor Staining Buffer Set from Thermo Fisher Scientific. Samples were acquired on a BD FACSCanto II or BD Symphony A1 flow cytometer, and data were analyzed using FlowJo software (v10.0.7).

High-dimensional T_reg_ cell analyses were conducted using *CytoML*, *flowWorkspace*, *flowCore*, and *Rtsne* packages in R. FlowJo workspaces and corresponding FCS files were imported, and the pre-gated live, single-cell CD4^+^ FoxP3^+^ cell population was extracted. Fluorescence channels corresponding to key markers were logicle-transformed and concatenated across all samples. Dimensionality reduction was performed using *t*-distributed stochastic neighbor embedding (t-SNE) on all gated events, and resulting plots were generated in *ggplot2*. Marker expression gradients were overlaid on the t-SNE maps using a shared color scale to visualize marker expression patterns.

### Ex vivo cell stimulation and functional assays

Non-parenchymal HMNCs were isolated as described above and cultured in complete RPMI medium (RPMI 1640 supplemented with 10% FBS, 0.1 mM MEM nonessential amino acids, 2 mM GlutaMAX-I, 10 mM HEPES, and 1 mM sodium pyruvate). Cells were seeded at a density of 0.5-1×10^6^ cells per well in U-bottom plates and stimulated with a combination of 50 ng/mL PMA and 500 ng/mL ionomycin, a combination of recombinant mouse IL-12 and IL-18 (5 ng/mL each), 2 nM 5-OP-RU, a 1:10 dilution of *E. coli* lysate, or with a 1:5 dilution of *K. pneumoniae* lysate. Bacterial lysates were prepared as previously described^72^. Stimulation with 5-OP-RU, *E. coli*, and *Klebsiella* was carried out in the presence of MR1-overexpressing M12 B cells as artificial antigen-presenting cells (APCs), which were generously provided by Dr. Jose Villadangos (University of Melbourne, Australia). Briefly, M12-MR1 cells were first pulsed with the aforementioned stimuli for 3 hours at 37°C, washed with PBS to remove excess antigen, and then co-cultured with HMNCs at a 5:1 APC-to-HMNC ratio. Co-cultures were maintained overnight except for PMA/ionomycin stimulation cultures, which lasted 4 hours. Brefeldin A was added to co-cultures at 10 µg/mL during the final 4 hours to inhibit protein secretion and allow for intracellular cytokine accumulation. Cells maintained in complete RPMI medium alone served as unstimulated controls. At the conclusion of co-cultures, cells were washed and stained for intracellular cytokine detection by flow cytometry.

### Bulk RNA-sequencing and analysis

HMNCs were isolated from *Mr1*^+/+^ and *Mr1*^-/-^ B6-MAIT^CAST^ mice receiving CCl_4_ 1, 12 or 26 days earlier as described above. Untreated mice were used as controls. Three mice were included per experimental group as biological replicates. Freshly isolated HMNCs were immediately processed for RNA extraction using the PureLink RNA Mini Kit (Thermo Fisher Scientific) following the manufacturer’s instructions. RNA samples were stored at -80°C until library preparation and sequencing. RNA quality and integrity were assessed using an Agilent 2100 Bioanalyzer, and all samples exhibited a 28S/18S rRNA ratio of ∼2.0.

Library preparation and sequencing were performed at the London Regional Genomics Centre (London, Canada) using an Illumina NextSeq High Output 75-cycle kit on an Illumina NextSeq platform, following the manufacturer’s mRNA-Seq workflow. Sequencing was performed to a depth of approximately 30 million paired-end reads per sample, providing sufficient coverage for transcriptome-wide differential expression analysis.

Raw sequencing reads were quality-checked with FastQC (v0.12.1), mapped to the *Mus musculus* reference genome (GRCm39) using STAR (v2.7.11b), and quantified with RSEM (v1.3.3). Downstream analyses were performed in R using *DESeq2* (v1.46.0). Variance-stabilizing transformation (VST) normalization and differential gene expression (DEG) analyses were conducted to compare CCl_4_-injected *Mr1*^+/+^ and *Mr1*^-/-^ mice at each time point, with samples from untreated animals serving as the baseline. Genes with an adjusted *p* < 0.05 and log_2_ fold change > 1 were considered significantly differentially expressed.

GSEA was performed using the Molecular Signatures Database (MSigDB) Hallmark^23^ gene sets to identify pathways significantly enriched for each comparison. Heatmaps were generated using expression values averaged across three biological replicates per group, normalized to baseline, and visualized for selected gene sets. A per-sample “T_reg_ cell score” was calculated by z-score normalizing each T_reg_-associated gene relative to its day 0 baseline. Values for each gene were then averaged, providing a single quantitative measure of T_reg_-related transcriptional activity per mouse.

### Interrogation of publicly available human single-cell RNA sequencing data

Publicly available single-cell RNA-seq data of CD45^+^ immune cells from human liver tissue, generated by Guilliams and colleagues, were obtained from CellXGene^44^. Samples with available single-cell data and corresponding fibrosis scoring were selected for analysis. Non-fibrotic samples were liver specimens without histopathological evidence of fibrosis (H06, H07, H10, H16, H22; 3 females, 2 males; ages 46–72 years), while fibrotic samples were defined as those with confirmed pathological fibrosis (H11, H13, H14, H18, H21, H23, H25; 6 females, 1 male; ages 28-77 years), as described in the original study **(Supplementary Table 1)**. Data processing and quality control were performed as described in the original publication, and preprocessed count matrices were further analyzed using *Seurat* (v5.1.0). Cell-type annotations from the original dataset were retained, with the exception of one cluster previously labeled as circulating effector memory T cells, which we reclassified as MAIT cells based on their high expression of *SLC4A10, KLRB1, ME1, LST1, NCR3, ZBTB16, TMIGD2,* and *IL23R*, consistent with a MAIT signature we previously defined^45^. The T_reg_ cell cluster was confirmed by the expression of *FOXP3, CD4, CTLA4* and *IKZF2*^73^.

Differential gene expression analyses for MAIT cells from healthy and fibrotic samples were carried out using Seurat’s *FindMarkers* function (Wilcoxon rank-sum test), with a false discovery rate (FDR)–adjusted *p* < 0.05 and log_2_ fold change > 0.1 being considered significant. Pathway enrichment analyses of differentially expressed genes were conducted using *clusterProfiler* and *fgsea* against the MSigDB Hallmark gene sets.

Dot plots were generated in Seurat to visualize the expression of gene sets associated with tissue residency, activation, exhaustion/dysfunction, cytokines, functional subsets, and tissue repair/fibrosis. Cell-cell communication analyses were performed using *CellChat* (v2.2.0) to infer ligand-receptor interactions between T_reg_ and MAIT cell clusters. Predicted signaling pathways and interaction networks were visualized according to the default workflow provided by the package.

### Statistical Analysis

Statistical comparisons were made using GraphPad Prism (v8.4.3) and JASP (v0.18.3.0). Normality was assessed using the Shapiro-Wilk test. Data were analyzed using two-tailed unpaired Student’s *t*-tests, Mann-Whitney *U*-tests, linear regression, two-way ANOVA with Sidak’s multiple comparisons test, or repeated-measures ANOVA with Sidak’s multiple comparisons test, as appropriate. A *p*-value < 0.05 was considered significant.

## ACKNOWLEDGMENTS

We thank Caroline O’Neil at Western University’s Molecular and Pathology Core Facility for her expert guidance and support with histological workflows.

This work was supported by the Canadian Institutes of Health Research (CIHR) (Project Grant #203791) and the Cancer Research Society (Operating Grant #1280925) to S.M.M.H. D.I.G. is supported by a National Health and Medical Research Council (NHMRC) investigator award (2008913). N.I.W. is supported by a Doctoral Research Award co-funded by the Cancer Research Society (CRS) and CIHR. V.S. received an Alexander Graham Bell Canada Graduate Scholarships – Master’s (CGS-M) from the Natural Sciences and Engineering Research Council of Canada (NSERC). A.M. is supported by a Postgraduate Scholarships – Doctoral program (PGS-D) from NSERC. A.S. is supported by an Ontario Graduate Scholarship.

Figures were created in BioRender, Wang, N. I. (2026).

## AUTHOR CONTRIBUTIONS

S.M.M.H. conceived the study, secured funding, contributed to experimental design and data interpretation, and supervised the project. N.I.W. contributed to experimental design, performed experiments, and analyzed the data. V.S. conducted the computational analyses of bulk RNA-Seq and single-cell RNA-Seq datasets. K.R., A.M., and A.S. contributed to data acquisition. B.A.K., S.Q.Z., A.R.M., and J.H. assisted in design, execution and/or interpretation of histological analyses. N.I.W. wrote the initial draft, which S.M.M.H. revised and edited.

## COMPETING INTERESTS STATEMENT

All authors declare no competing interests.

---

**Supplementary Fig. 1.**
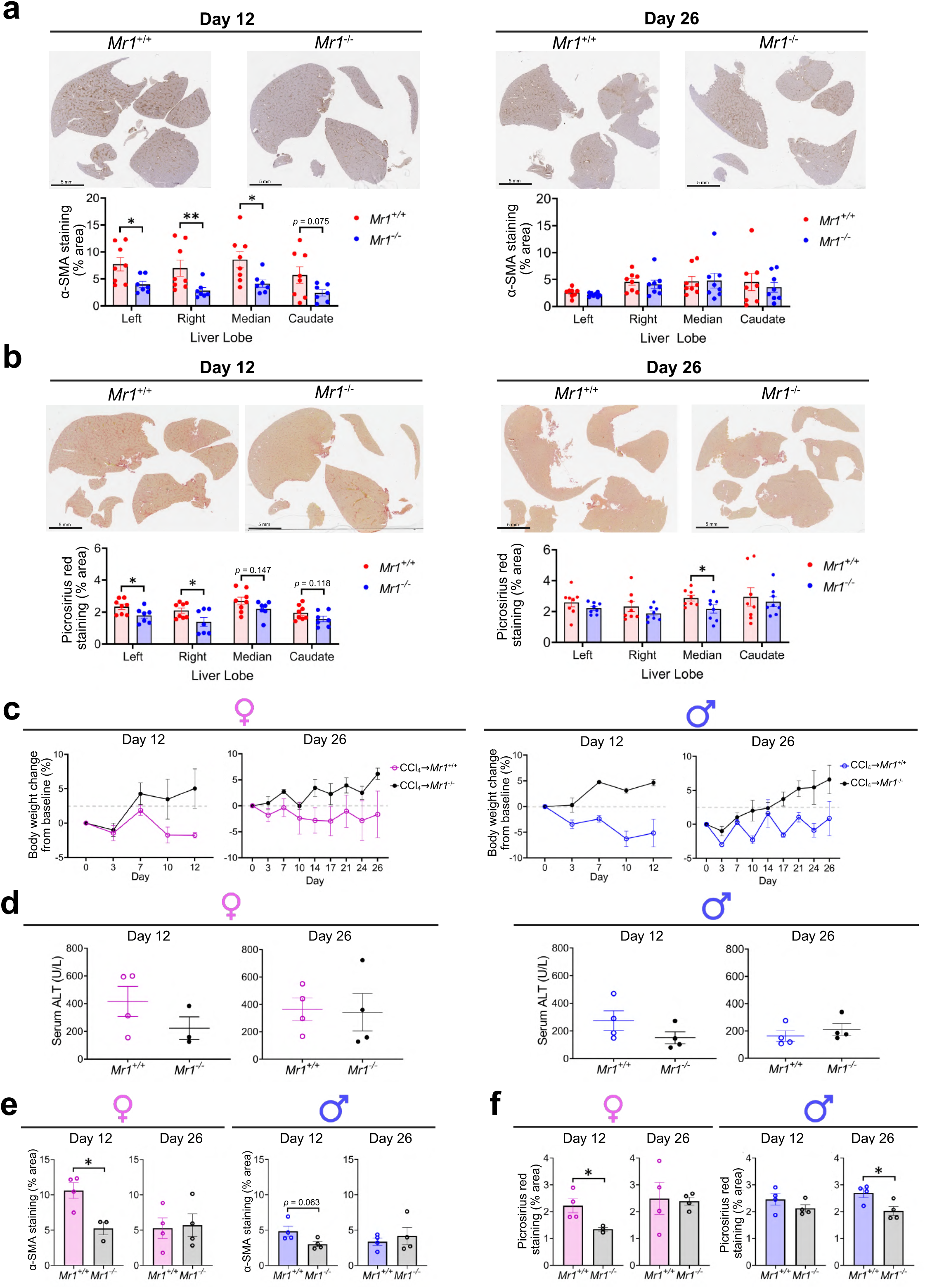
Sex-stratified and whole-liver histological analyses of fibrosis and injury. Whole-slide imaging of liver sections from *Mr1^+/+^* and *Mr1^-/-^* mice was performed following α-SMA (a) and Picrosirius Red (b) staining to assess global fibrotic area and collagen deposition, respectively, and quantified across individual liver lobes (left, right, median, and caudate). Body weight change from baseline (c) and serum ALT levels (d) were measured in CCl_4_-treated *Mr1^+/+^* and *Mr1^-/-^* mice in a sex-stratified manner at days 12 and 26. Fibrosis was further evaluated by quantifying α-SMA⁺ area (e) and Picrosirius Red⁺ area (f) in female and male mice at both time points. Data are pooled from 4 (a-b) and 2 (c-f) independent experiments. Data are presented as mean ± SEM. Statistical analyses were performed using unpaired Student’s *t* tests, Mann-Whitney *U* tests, or repeated-measures ANOVAs. * and ** denote differences with *p* < 0.05 and 0.01, respectively.

**Supplementary Fig. 2.**
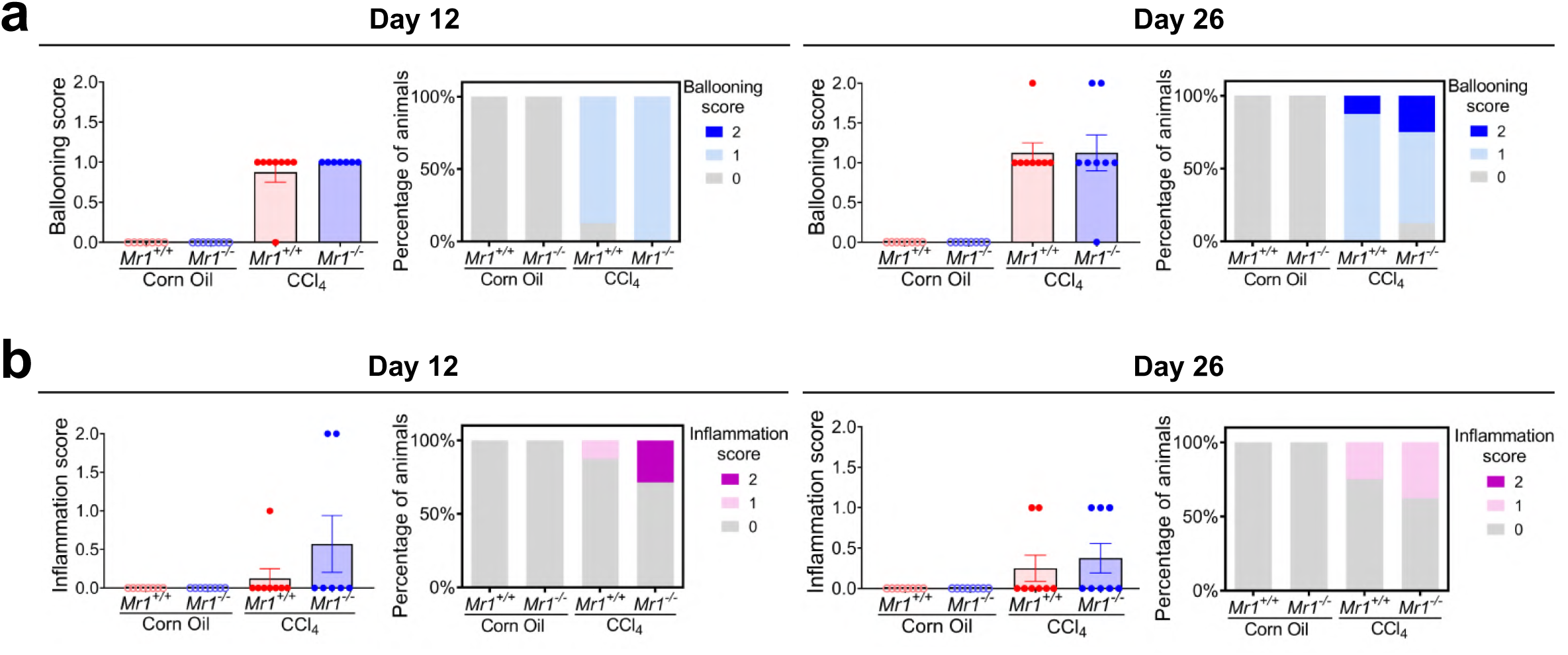
Histopathological assessment of hepatocellular ballooning and inflammation. Hepatocellular ballooning (a) and inflammation (b) were scored by board-certified pathologists using H&E-stained liver sections from *Mr1^+/+^* and *Mr1^-/-^* mice and are presented as raw scores and the proportion of animals within each score category (0, 1, 2). Data are pooled from 4 independent experiments. Data are presented as mean ± SEM and stacked bar graphs. Statistical analyses were performed using two-way ANOVAs.

**Supplementary Fig. 3.**
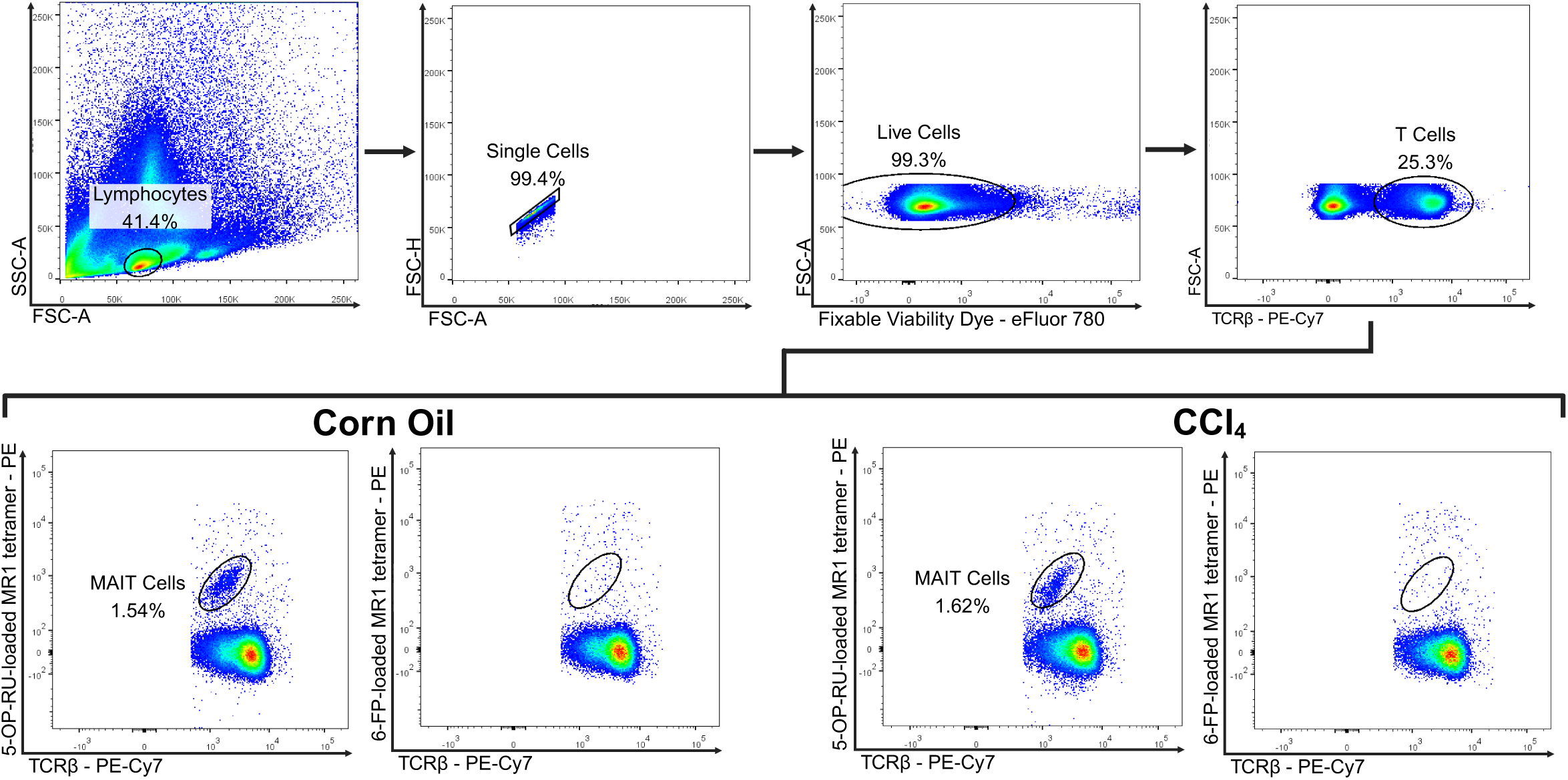
Flow cytometry gating strategy for MAIT cell identification. Cells were sequentially gated on lymphocytes, singlets, live cells, and TCRβ⁺ cells, followed by identification of MAIT cells using 5-OP-RU-loaded MR1 tetramer. 6-FP-loaded MR1 tetramer was used as a negative control.

**Supplementary Fig. 4.**
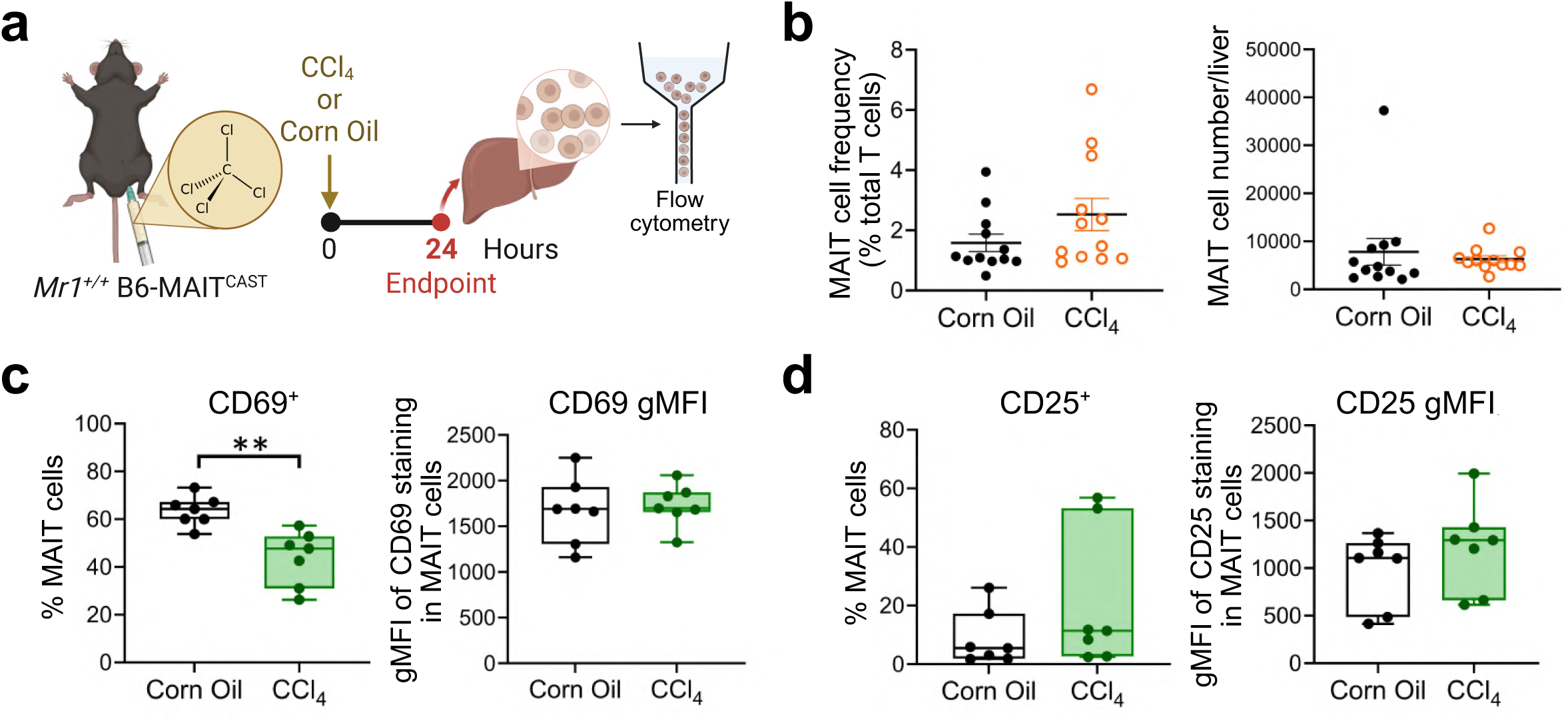
MAIT cell activation in an acute model of liver injury. *Mr1^+/+^* mice were administered CCl_4_ or corn oil and analyzed by flow cytometry 24 hours later (a). The frequency of hepatic MAIT cells among T cells and absolute MAIT cell numbers were quantified following treatment (b). CD69 (c) and CD25 (d) expression on MAIT cells were assessed by measuring the frequency of positive cells and gMFI. Data are pooled from 5 (b) and 3 (c-d) independent experiments. Data are presented as mean ± SEM (b) and box-and-whisker plots showing median, IQR, and min-max (c-d). Statistical analyses were performed using Student’s *t* tests or Mann-Whitney *U* tests. ** denotes differences with *p* < 0.01.

**Supplementary Fig. 5.**
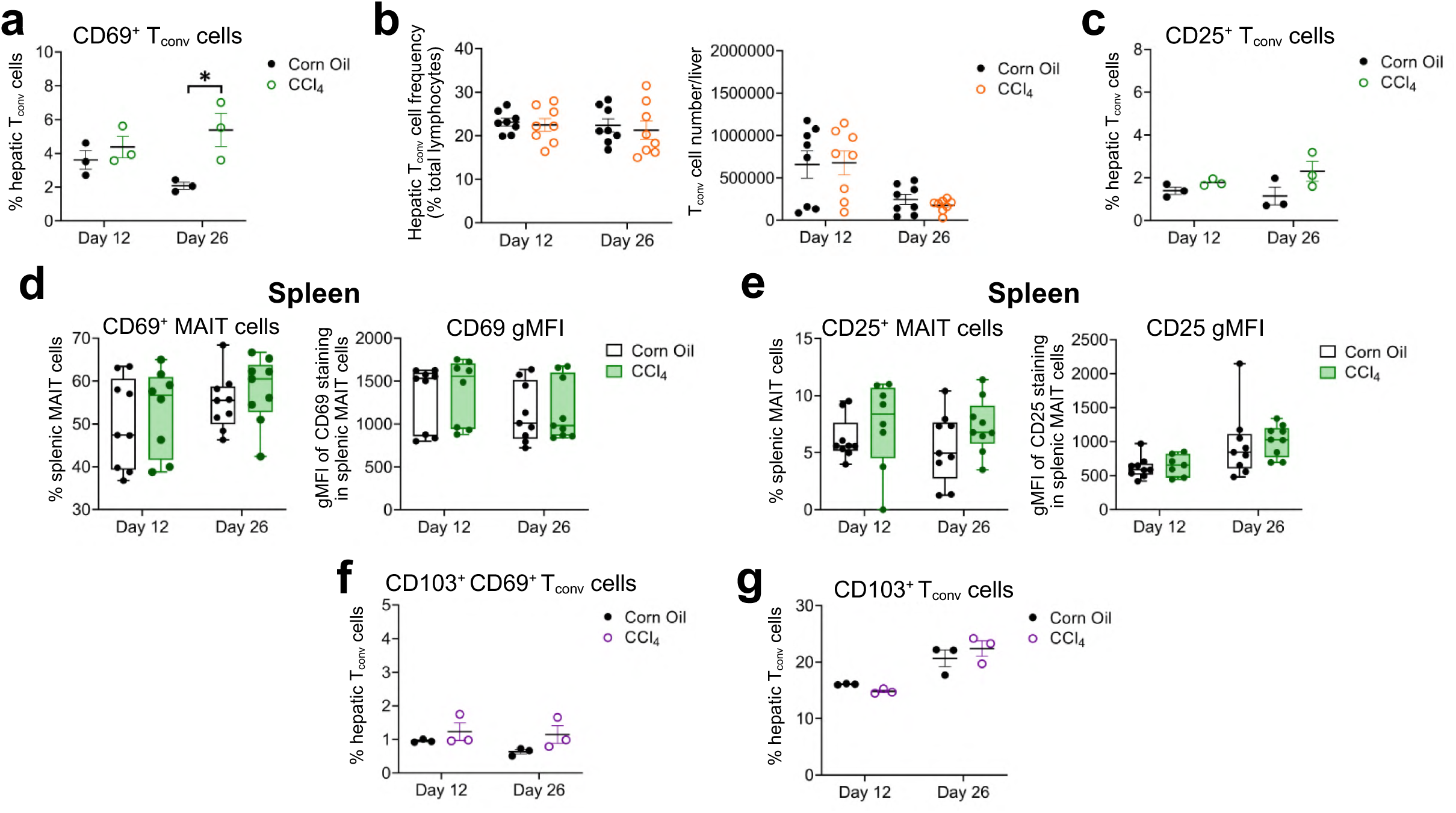
Activation and tissue residency of hepatic T_conv_ cells and splenic MAIT cells. Hepatic T_conv_ (5-OP-RU-loaded mouse MR1 tetramer^−^, PBS-57-loaded mouse CD1d tetramer^−^, TCRβ^+^) cells were characterized by CD69^+^ frequency (a), overall cell frequency and absolute number (b), and CD25^+^ frequency (c) at days 12 and 26 following corn oil or CCl_4_ treatment. Activation of splenic MAIT cells was assessed by measuring CD69 (d) and CD25 (e) expression as the frequency of positive cells and gMFI of each marker. Tissue residency- associated phenotypes of hepatic T_conv_ cells were examined by quantifying the CD103^+^ CD69^+^ (f) and CD103^+^ (g) populations at both time points. Data in (b, d-e) are pooled from 3 independent experiments. Data are presented as mean ± SEM (a-c, f-g) and box-and-whisker plots showing median, IQR, and min-max (d-e). Statistical analyses were performed using Student’s *t* tests or Mann-Whitney *U* tests. * denotes differences with *p* < 0.05.

**Supplementary Fig. 6.**
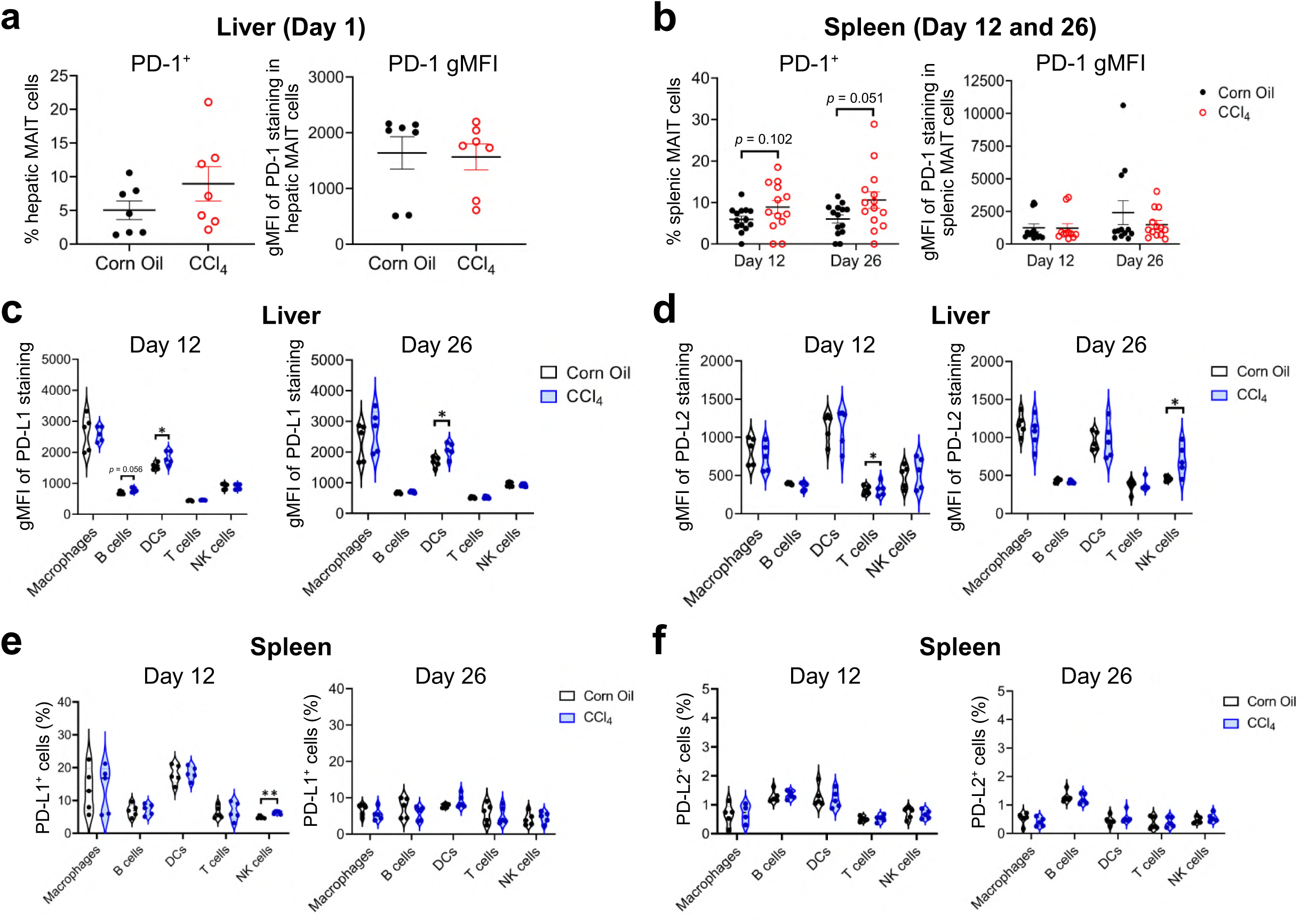
PD-1 and PD-1 ligand expression on hepatic and splenic immune cells. PD-1 expression on MAIT cells was assessed cytofluorometrically in the liver at 24 hours (a) and in the spleen at days 12 and 26 (b) and is presented as the frequency of PD-1^+^ cells and PD-1 gMFI. gMFI of PD-L1 (c) and PD-L2 (d) staining on hepatic immune cell subsets (macrophages, B cells, dendritic cells, T cells, and NK cells) was evaluated at days 12 and 26. In the spleen, PD-L1^+^ (e) and PD-L2^+^ (f) immune cell populations were quantified across the same cell subsets and time points. Data are pooled from 3 (a), 5 (b), and 2 (c-f) independent experiments. Data are presented as mean ± SEM (a-b) and violin plots showing distribution, median, and quartiles (c-f). Statistical analyses were performed using Student’s *t* tests or Mann-Whitney *U* tests. * and ** denote differences with *p* < 0.05 and 0.01, respectively.

**Supplementary Fig. 7.**
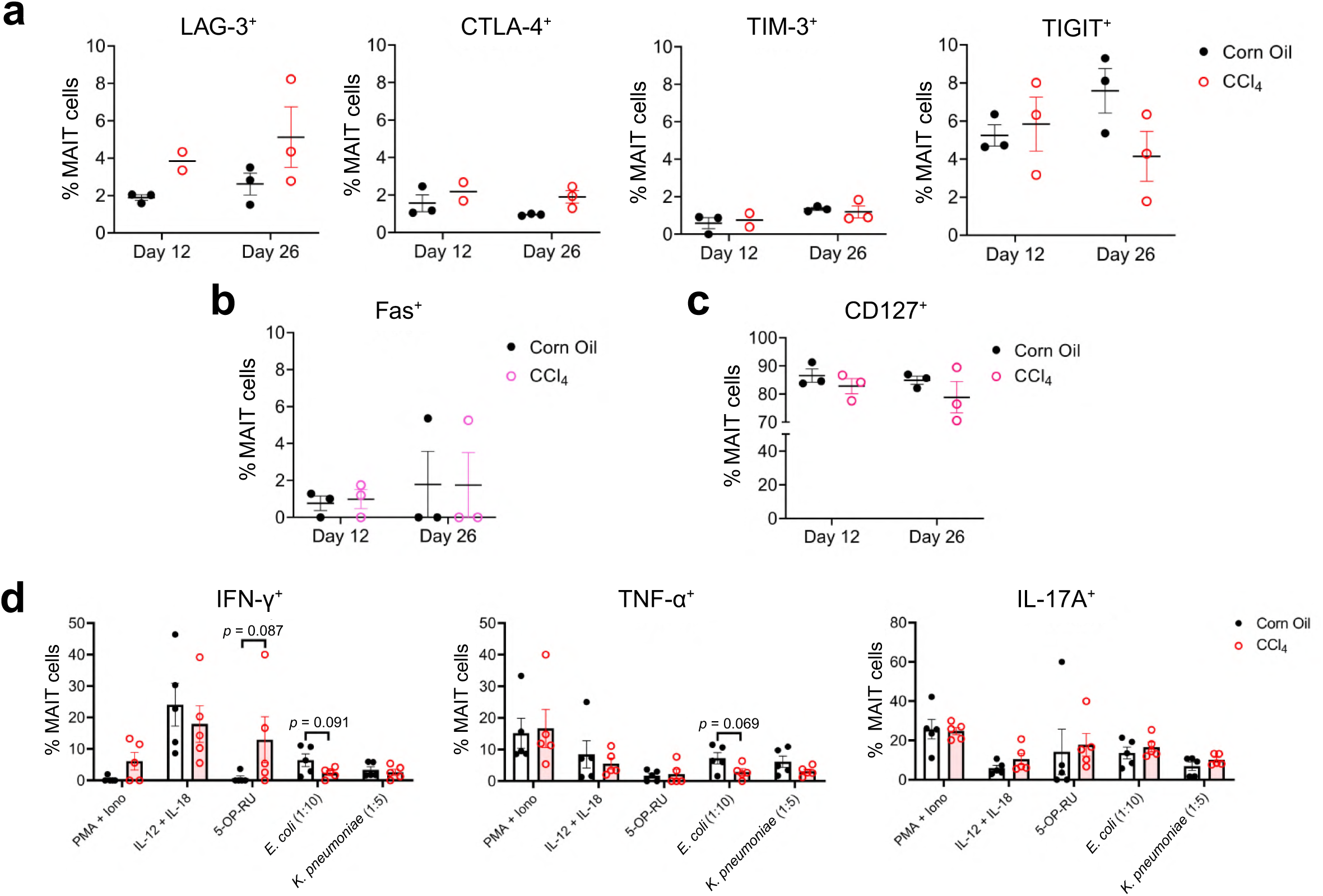
Additional characterization of hepatic MAIT cell exhaustion. Expression of exhaustion and co-inhibitory markers (LAG-3, CTLA-4, TIM-3, and TIGIT) on hepatic MAIT cells was evaluated at days 12 and 26 (a). Expression of Fas (b) and CD127 (c) on hepatic MAIT cells was also assessed at the same time points. Cytokine production by hepatic MAIT cells was measured at day 12 following *ex vivo* stimulation with PMA and ionomycin, IL-12 and IL-18, 5-OP-RU, *E. coli* lysate, or *K. pneumoniae* lysate, and is presented as the frequency of IFN-γ^+^, TNF^+^, and IL-17A^+^ cells (d). Data in (d) are pooled from 2 independent experiments. Data are presented as mean ± SEM. Statistical analyses were performed using Student’s *t* tests or Mann-Whitney *U* tests.

**Supplementary Fig. 8.**
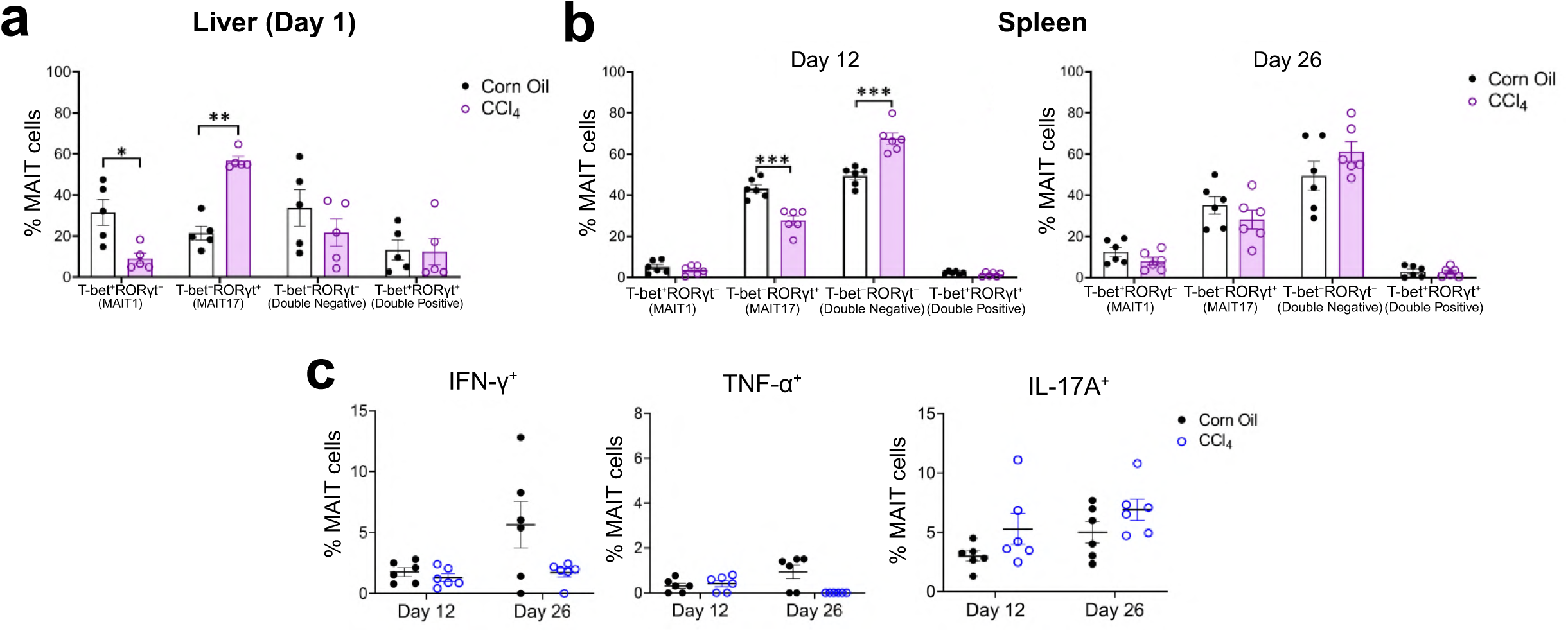
MAIT1 and MAIT17 polarization in acute liver injury and chronic splenic responses. The frequency of hepatic T-bet^+^RORγt^−^ (MAIT1), T-bet^−^RORγt^+^ (MAIT17), T-bet^−^RORγt^−^ (Double Negative), and T-bet^+^RORγt^+^ (Double Positive) MAIT cells was quantified 24 hours following corn oil and CCl_4_ treatment. The same MAIT cell subsets were assessed in the spleen at days 12 and 26 (b), alongside baseline cytokine production, presented as the frequency of IFN-γ^+^, TNF^+^, and IL-17A^+^ MAIT cells (c). Data in are pooled from 2 independent experiments. Data are presented as mean ± SEM. Statistical analyses were performed using unpaired Student’s *t* tests or Mann-Whitney *U* tests. *, **, and *** denote differences with *p* < 0.05, 0.01, and 0.001, respectively.

**Supplementary Fig. 9.**
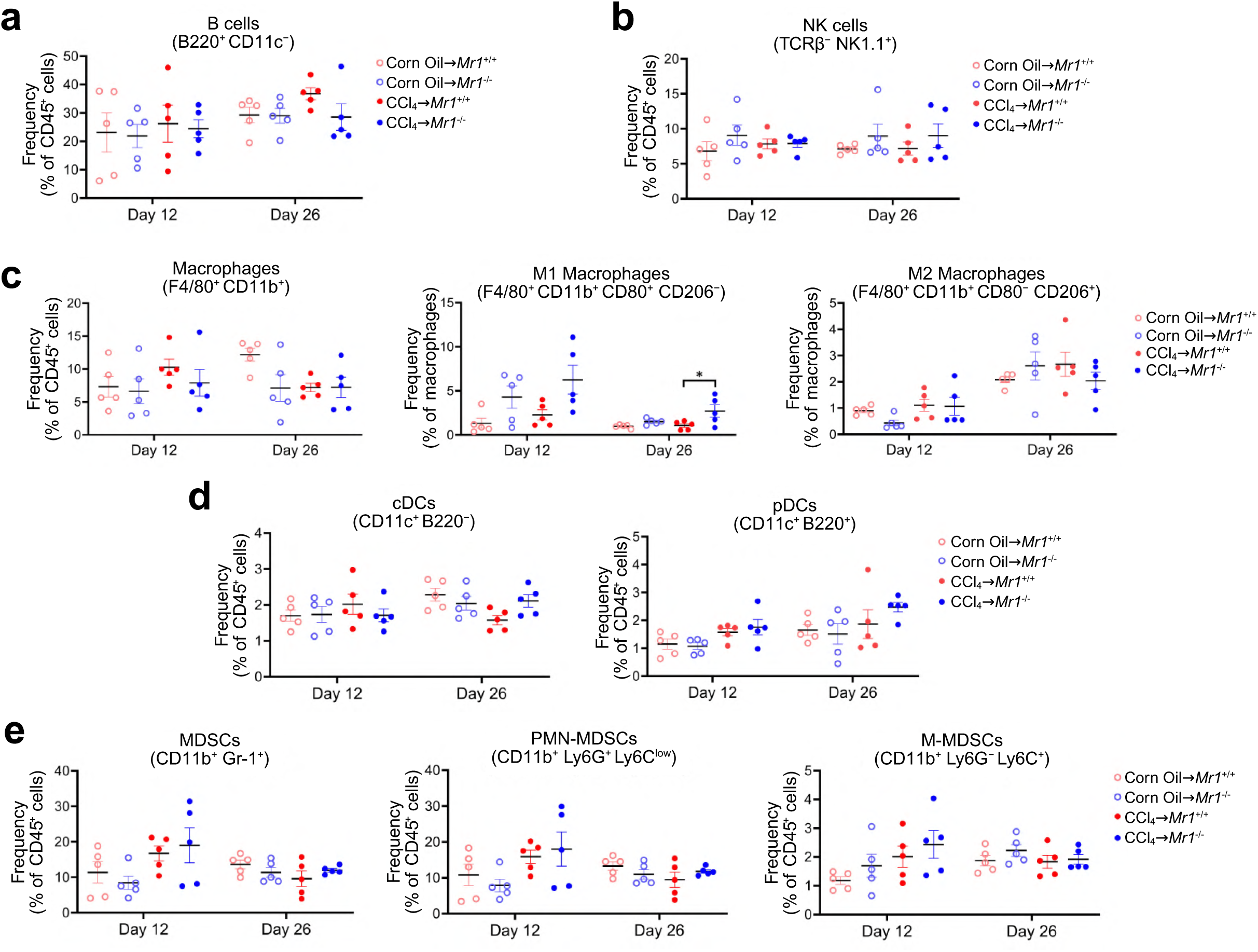
Hepatic immune landscape in *Mr1^+/+^* and *Mr1^+/+^* mice during CCl_4_-induced injury. Hepatic immune cell populations, including B cells (a), NK cells (b), macrophages (total, M1, and M2) (c), DCs (conventional DCs and plasmacytoid DCs) (d), and MDSCs (total, polymorphonuclear MDSCs, and monocytic MDSCs) (e), were quantified flow cytometrically in corn oil- or CCl_4_-treated *Mr1^+/+^* and *Mr1^-/-^* mice at the indicated time points. Data in are pooled from 2 independent experiments. Data are presented as mean ± SEM. Statistical analyses were performed using two-way ANOVAs. * denotes differences with *p* < 0.05.

**Supplementary Fig. 10.**
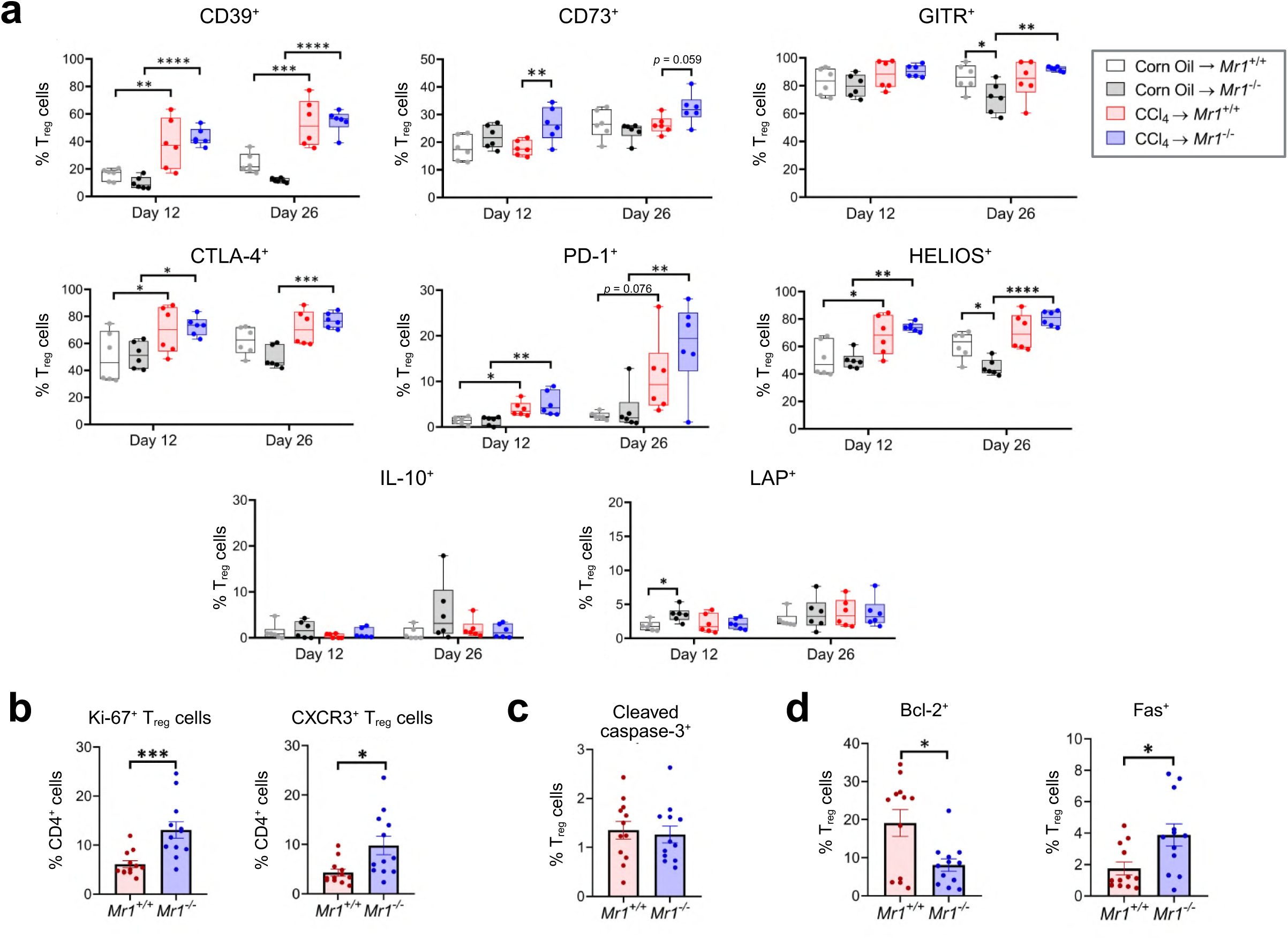
Additional characterization of hepatic T_reg_ cells. Expression of CD39, CD73, GITR, CTLA-4, PD-1, HELIOS, IL-10, and LAP on hepatic T_reg_ cells was evaluated by flow cytometry in corn oil- or CCl_4_-treated *Mr1^+/+^* and *Mr1^-/-^* mice at days 12 and 26 (a). Proliferation and trafficking were further assessed by quantifying Ki-67^+^ and CXCR3^+^ T_reg_ cells as a fraction of all CD4^+^ T cells (b). Apoptosis and survival markers were examined by measuring cleaved caspase-3^+^ (c), Bcl-2^+^, and Fas^+^ (d) T_reg_ cells. Data are pooled from 2 independent experiments. Data are presented as mean ± SEM. Statistical analyses were performed using two-way ANOVAs (a), Student’s *t* tests, or Mann-Whitney *U* tests (b-d). *, **, ***, and **** denote differences with *p* < 0.05, 0.01, 0.001, and 0.0001, respectively.

**Supplementary Fig. 11.**
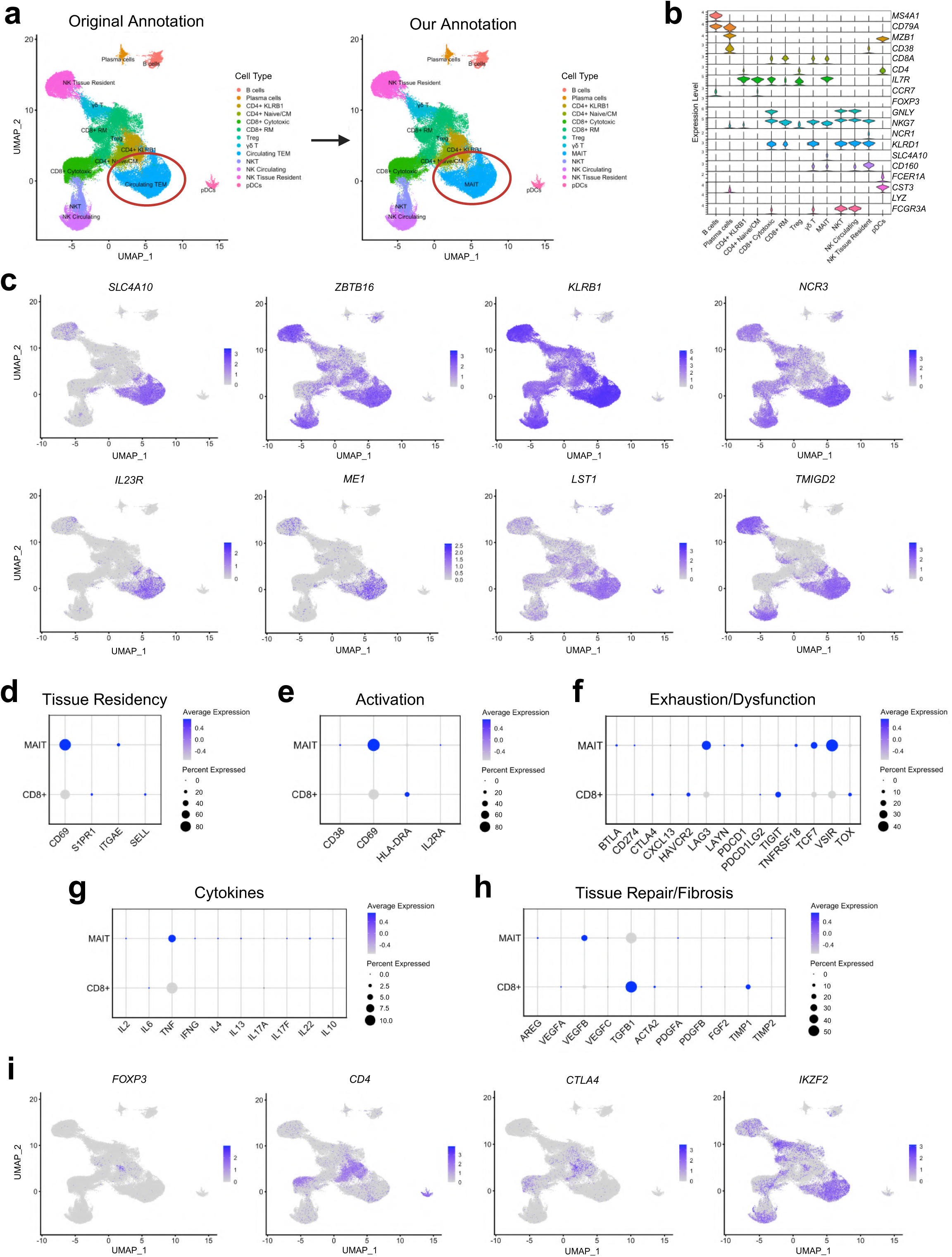
Re-annotation and transcriptional comparison of MAIT cells and CD8^+^ T cells in human liver single-cell RNA-seq data. UMAP visualization of CD45^+^ immune cells with the original (left) and our updated (right) annotations highlights reclassification of the circulating effector memory T cell (T_EM_) cluster as MAIT cells based on marker gene expression (a). Expression of canonical lineage-defining genes across annotated immune populations supports the revised cluster identities (b). Feature plots of MAIT cell-associated genes (*SLC4A10*, *ZBTB16*, *KLRB1*, *NCR3*, *IL23R*, *ME1*, *LST1*, and *TMIGD2*) further validate this re-annotation (c). Dot plots comparing MAIT cells and cytotoxic CD8^+^ T cells illustrate functional gene signatures related to tissue residency (d), activation (e), exhaustion/dysfunction (f), cytokine production (g), and tissue repair/fibrosis (h). Feature plots of T_reg_-associated genes (*FOXP3*, *CD4*, *CTLA4*, and *IKZF2*) are also shown (i).

**Supplementary Table 1.** Clinical characteristics of non-fibrotic and fibrotic samples used for single-cell RNA-seq analysis.

| Analysis Group | Guilliams *et al.* Sample ID | Gender | Age (years) | Steatosis | Fibrosis |
| --- | --- | --- | --- | --- | --- |
| Non-fibrotic | H06 | Male | 72 | No | No |
|  | H07 | Female | 46 | No | No |
|  | H10 | Male | 55 | No | No |
|  | H16 | Female | 66 | <5% steatosis | No |
|  | H22 | Female | 55 | No | No |
| Fibrotic | H11 | Female | 53 | Mild (20%) | Clear pericellular fibrosis |
|  | H13 | Male | 73 | 5% steatosis | Clear pericellular fibrosis |
|  | H14 | Female | 75 | >5% steatosis | Very mild pericellular fibrosis |
|  | H18 | Female | 28 | No | Mild pericellular fibrosis |
|  | H21 | Female | 77 | <5% steatosis | Cirrhosis |
|  | H23 | Female | 75 | No | 5-10% fibrosis |
|  | H25 | Female | 49 | 5-10% steatosis | Minimal pericellular fibrosis |

**Supplemental Table 2.** Antibodies, isotype controls, tetramers and viability dyes used for flow cytometry.

| **Antibodies** | **Clone** | **Company** |
| --- | --- | --- |
| PE-Cyanine7-conjugated anti-mouse TCRβ | H57-597 | Thermo Fisher Scientific |
| Alexa Fluor 700-conjugated anti-mouse TCRβ | H57-597 | Thermo Fisher Scientific |
| PE-eFluor 610-conjugated anti-mouse CD25 | PC61.5 | Thermo Fisher Scientific |
| PE-Cyanine5.5-conjugated anti-mouse CD25 | PC61.5 | Thermo Fisher Scientific |
| PerCP-Cyanine5.5-conjugated anti-mouse CD69 | H1.2F3 | Thermo Fisher Scientific |
| APC-conjugated anti-mouse CTLA-4 | UC10-4B9 | Thermo Fisher Scientific |
| PE-eFluor 610-conjugated anti-mouse CTLA-4 | UC10-4B9 | Thermo Fisher Scientific |
| APC-conjugated anti-mouse TIM-3 | 8B.2C12 | Thermo Fisher Scientific |
| PerCP-eFluor 710-conjugated anti-mouse LAG-3 | C9B7W | Thermo Fisher Scientific |
| PE-eFluor 610-conjugated anti-mouse PD-1 | J43 | Thermo Fisher Scientific |
| PE-Cyanine7-conjugated anti-mouse PD-1 | J43 | Thermo Fisher Scientific |
| eFluor 506-conjugated anti-mouse PD-1 | J43 | Thermo Fisher Scientific |
| PE-conjugated anti-mouse CD103 | 2E7 | Thermo Fisher Scientific |
| Alexa Fluor 700-conjugated anti-mouse CD103 | 2E7 | Thermo Fisher Scientific |
| PerCP-eFluor 710-conjugated anti-mouse TIGIT | GIGD7 | Thermo Fisher Scientific |
| PE-conjugated anti-mouse BTLA | 8F4 | Thermo Fisher Scientific |
| PE-conjugated anti-mouse T-bet | eBio4B10 | Thermo Fisher Scientific |
| PE-eFluor 610-conjugated anti-mouse GATA-3 | TWAJ | Thermo Fisher Scientific |
| PerCP-eFluor 710-conjugated anti-mouse RORγt | B2D | Thermo Fisher Scientific |
| APC-conjugated anti-mouse TNF-α | MP6-XT22 | Thermo Fisher Scientific |
| PerCP-Cyanine5.5-conjugated anti-mouse IFN-γ | XMG1.2 | Thermo Fisher Scientific |
| PE-eFluor 610-conjugated anti-mouse IL-17A | eBio1787 | Thermo Fisher Scientific |
| APC-conjugated anti-mouse CD122 | TM-b1 | Thermo Fisher Scientific |
| PE-eFluor 610-conjugated anti-mouse CD62L | MEL-14 | Thermo Fisher Scientific |
| PerCP-eFluor 710-conjugated anti-mouse Fas | 15A7 | Thermo Fisher Scientific |
| PE-conjugated anti-mouse Fas | 15A7 | Thermo Fisher Scientific |
| APC-conjugated anti-mouse CD127 | SB/199 | Thermo Fisher Scientific |
| Alexa Fluor 700-conjugated anti-mouse CD45 | 30-F11 | Thermo Fisher Scientific |
| PE-Cyanine7-conjugated anti-mouse CD45 | 30-F11 | Thermo Fisher Scientific |
| PerCP-eFluor 710-conjugated anti-mouse PD-L1 | MIH5 | Thermo Fisher Scientific |
| PE-conjugated anti-mouse PD-L2 | 122 | Thermo Fisher Scientific |
| APC-conjugated anti-mouse F4/80 | BM8 | Thermo Fisher Scientific |
| PE-eFluor 610-conjugated anti-mouse CD11b | M1/70 | Thermo Fisher Scientific |
| Alexa Fluor 700-conjugated anti-mouse CD11b | M1/70 | Thermo Fisher Scientific |
| APC-conjugated anti-mouse B220 | RA3-6B2 | Thermo Fisher Scientific |
| PE-Cyanine7-conjugated anti-mouse NK1.1 | PK136 | Thermo Fisher Scientific |
| PE-eFluor 610-conjugated anti-mouse CD11c | N418 | Thermo Fisher Scientific |
| PE-conjugated anti-mouse Tox | TXRX10 | Thermo Fisher Scientific |
| PE-conjugated anti-mouse CD80 | 16-10A1 | Thermo Fisher Scientific |
| PerCP-eFluor 710-conjugated anti-mouse CD206 | MR6F3 | Thermo Fisher Scientific |
| PE-Cyanine7-conjugated anti-mouse Gr-1 | RB6-8C5 | Thermo Fisher Scientific |
| PerCP-Cyanine5.5-conjugated anti-mouse Ly-6C | HK1.4 | BioLegend |
| APC-conjugated anti-mouse Ly-6G | 1AB-Ly6g | Thermo Fisher Scientific |
| PE-eFluor 610-conjugated anti-mouse CD4 | GK1.5 | Thermo Fisher Scientific |
| PE-conjugated anti-mouse CD4 | GK1.5 | Thermo Fisher Scientific |
| PE-Cy5-conjugated anti-mouse CD4 | GK1.5 | Thermo Fisher Scientific |
| Alexa Fluor 700-conjugated anti-mouse CD4 | GK1.5 | Thermo Fisher Scientific |
| PerCP-Cyanine5.5-conjugated anti-mouse FoxP3 | FJK-16s | Thermo Fisher Scientific |
| PerCP-eFluor 710-conjugated anti-mouse FoxP3 | FJK-16s | Thermo Fisher Scientific |
| Super Bright 702-conjugated anti-mouse CD39 | 24DMS1 | Thermo Fisher Scientific |
| eFluor 450-conjugated anti-mouse CD73 | eBioTY/11.8 | Thermo Fisher Scientific |
| PE-Cyanine7-conjugated anti-mouse LAP | TW7-16B4 | Thermo Fisher Scientific |
| Super Bright 600-conjugated anti-mouse GITR | DTA-1 | Thermo Fisher Scientific |
| PE-conjugated anti-mouse HELIOS | 22F6 | Thermo Fisher Scientific |
| APC-conjugated anti-mouse IL-10 | JES5-16E3 | Thermo Fisher Scientific |
| PE-eFluor 610-conjugated anti-mouse Ki-67 | SolA15 | Thermo Fisher Scientific |
| PE-conjugated anti-mouse cleaved caspase-3 | C92-605.rMAb | BD Biosciences |
| PE-Cyanine7-conjugated anti-mouse Bcl-2 | 10C4 | Thermo Fisher Scientific |
| PerCP-eFluor 710-conjugated anti-mouse CXCR3 | CXCR3-173 | Thermo Fisher Scientific |

| **Isotype controls** | **Clone** | **Company** |
| --- | --- | --- |
| PE-eFluor 610-conjugated Rat IgG1κ | eBRG1 | Thermo Fisher Scientific |
| PerCP-Cyanine5.5-conjugated Armenian Hamster IgG | eBio299Arm | Thermo Fisher Scientific |
| APC-conjugated Armenian Hamster IgG | eBio299Arm | Thermo Fisher Scientific |
| Alexa Fluor 700-conjugated Armenian Hamster IgG | eBio299Arm | Thermo Fisher Scientific |
| APC-conjugated mouse IgG1κ | P3.6.2.8.1 | Thermo Fisher Scientific |
| PerCP-eFluor 710-conjugated Rat IgG1κ | eBRG1 | Thermo Fisher Scientific |
| PE-eFluor 610-conjugated Armenian Hamster IgG | eBio299Arm | Thermo Fisher Scientific |
| PE-conjugated Armenian Hamster IgG | eBio299Arm | Thermo Fisher Scientific |
| PerCP-eFluor 710-conjugated Rat IgG2ak | eBR2a | Thermo Fisher Scientific |
| PE-conjugated mouse IgG1κ | P3.6.2.8.1 | Thermo Fisher Scientific |
| PE-eFluor 610-conjugated Rat IgG2ak | eBR2a | Thermo Fisher Scientific |
| PE-Cyanine7-conjugated Rat IgG2bκ | eB149/10H5 | Thermo Fisher Scientific |
| PerCP-eFluor 710-conjugated Rat IgG2bκ | eB149/10H5 | Thermo Fisher Scientific |
| PE-eFluor 610-conjugated Rat IgG2bκ | eB149/10H5 | Thermo Fisher Scientific |
| APC-conjugated Rat IgG2bκ | eB149/10H5 | Thermo Fisher Scientific |
| PerCP-Cyanine5.5-conjugated Rat IgG2aκ | eBR2a | Thermo Fisher Scientific |
| PE-Cyanine7-conjugated Armenian Hamster IgG | eBio299Arm | Thermo Fisher Scientific |
| APC-conjugated Rat IgG1κ | eBRG1 | Thermo Fisher Scientific |
| PerCP-Cyanine5.5-conjugated Rat IgG1κ | eBRG1 | Thermo Fisher Scientific |
| PerCP-eFluor 710-conjugated mouse IgG1κ | P3.6.2.8.1 | Thermo Fisher Scientific |
| PE-conjugated Rat IgG2aκ | eBR2a | Thermo Fisher Scientific |
| Super Bright 702-conjugated Rat IgG2bκ | eB149/10H5 | Thermo Fisher Scientific |
| eFluor 450-conjugated Rat IgG1κ | eBRG1 | Thermo Fisher Scientific |
| PE-Cyanine7-conjugated mouse IgG1κ | P3.6.2.8.1 | Thermo Fisher Scientific |
| Super Bright 600-conjugated Rat IgG2bκ | eB149/10H5 | Thermo Fisher Scientific |
| eFluor 506-conjugated Armenian Hamster IgG | eBio299Arm | Thermo Fisher Scientific |

| **Tetramers** | **Source** |
| --- | --- |
| PE-conjugated 5-OP-RU-loaded mouse MR1 tetramer | NIH Tetramer Core Facility |
| PE-conjugated 6-FP-loaded mouse MR1 tetramer | NIH Tetramer Core Facility |
| APC-conjugated 5-OP-RU-loaded mouse MR1 tetramer | NIH Tetramer Core Facility |
| APC-conjugated 6-FP-loaded mouse MR1 tetramer | NIH Tetramer Core Facility |
| PE-conjugated PBS-57-loaded mouse CD1d tetramer | NIH Tetramer Core Facility |
| PE-conjugated unloaded mouse CD1d tetramer | NIH Tetramer Core Facility |
| APC-conjugated PBS-57-loaded mouse CD1d tetramer | NIH Tetramer Core Facility |
| APC-conjugated unloaded mouse CD1d tetramer | NIH Tetramer Core Facility |

| **Viability Dyes** | **Company** |
| --- | --- |
| Fixable viability dye-eFluor780 | Thermo Fisher Scientific |

## Notes

### Competing Interest Statement

The authors have declared no competing interest.

## REFERENCES

1. Zamani, M. et al. Global prevalence of advanced liver fibrosis and cirrhosis in the general population: A systematic review and meta-analysis. Clin Gastroenterol Hepatol 23, 1123–1134 (2025).

2. Devarbhavi, H. et al. Global burden of liver disease: 2023 update. J Hepatol 79, 516–537 (2023).

3. Kisseleva, T. & Brenner, D. Molecular and cellular mechanisms of liver fibrosis and its regression. Nat Rev Gastroenterol Hepatol 18, 151–166 (2021).

4. Roehlen, N., Crouchet, E. & Baumert, T. F. Liver fibrosis: Mechanistic concepts and therapeutic perspectives. Cells 9, 875 (2020).

5. Kurioka, A., Walker, L. J., Klenerman, P. & Willberg, C. B. MAIT cells: new guardians of the liver. Clin Transl Immunology 5, e98 (2016).

6. Rahimpour, A. et al. Identification of phenotypically and functionally heterogeneous mouse mucosal-associated invariant T cells using MR1 tetramers. J Exp Med 212, 1095–1108 (2015).

7. Kjer-Nielsen, L. et al. MR1 presents microbial vitamin B metabolites to MAIT cells. Nature 491, 717–723 (2012).

8. Ussher, J. E. et al. CD161^++^ CD8^+^ T cells, including the MAIT cell subset, are specifically activated by IL-12+IL-18 in a TCR-independent manner. Eur J Immunol 44, 195–203 (2014).

9. Nel, I., Bertrand, L., Toubal, A. & Lehuen, A. MAIT cells, guardians of skin and mucosa? Mucosal Immunol 14, 803–814 (2021).

10. Zhang, M. & Zhang, S. T cells in fibrosis and fibrotic diseases. Front Immunol 11, 1142 (2020).

11. Hinks, T. S. C. et al. Activation and in vivo evolution of the MAIT cell transcriptome in mice and humans reveals tissue repair functionality. Cell Rep 28, 3249–3262.e5 (2019).

12. Constantinides, M. G. et al. MAIT cells are imprinted by the microbiota in early life and promote tissue repair. Science 366, eaax6624 (2019).

13. du Halgouet, A. et al. Role of MR1-driven signals and amphiregulin on the recruitment and repair function of MAIT cells during skin wound healing. Immunity 56, 78–92.e6 (2023).

14. Hegde, P. et al. Mucosal-associated invariant T cells are a profibrogenic immune cell population in the liver. Nat Commun 9, 2146 (2018).

15. Böttcher, K. et al. MAIT cells are chronically activated in patients with autoimmune liver disease and promote profibrogenic hepatic stellate cell activation. Hepatology 68, 172–186 (2018).

16. Mabire, M. et al. MAIT cell inhibition promotes liver fibrosis regression via macrophage phenotype reprogramming. Nat Commun 14, 1830 (2023).

17. Jiang, X. et al. MAIT cells ameliorate liver fibrosis by enhancing the cytotoxicity of NK cells in cholestatic murine models. Liver Int 42, 2743–2758 (2022).

18. Cui, Y. et al. Mucosal-associated invariant T cell–rich congenic mouse strain allows functional evaluation. J Clin Invest 125, 4171–4185 (2015).

19. Habaz, I. A. et al. MAIT cells promote cancer progression and regulatory T cell accumulation in bladder tumor microenvironment. J Immunother Cancer 13, e012496 (2025).

20. Rashu, R. et al. Targeting the MR1-MAIT cell axis improves vaccine efficacy and affords protection against viral pathogens. PLOS Pathogens 19, e1011485 (2023).

21. Starkel, P. & Leclercq, I. A. Animal models for the study of hepatic fibrosis. Best Pract Res Clin Gastroenterol 25, 319–333 (2011).

22. Scholten, D., Trebicka, J., Liedtke, C. & Weiskirchen, R. The carbon tetrachloride model in mice. Lab Anim 49, 4–11 (2015).

23. Liberzon, A. et al. The Molecular Signatures Database (MSigDB) hallmark gene set collection. Cell Syst 1, 417–425 (2015).

24. Gao, R. et al. Comprehensive analysis of endoplasmic reticulum-related and secretome gene expression profiles in the progression of non-alcoholic fatty liver disease. Front Endocrinol (Lausanne) 13, 967016 (2022).

25. Oshi, M. et al. The E2F Pathway Score as a Predictive Biomarker of Response to Neoadjuvant Therapy in ER+/HER2- Breast Cancer. Cells 9, 1643 (2020).

26. Liao, R. et al. E2F transcription factor 1 (E2F1) promotes the transforming growth factor TGF-β1 induced human cardiac fibroblasts differentiation through promoting the transcription of CCNE2 gene. Bioengineered 12, 6869–6877 (2021).

27. Zacarías-Fluck, M. F., Soucek, L. & Whitfield, J. R. MYC: there is more to it than cancer. Front Cell Dev Biol 12, 1342872 (2024).

28. Woodcock, H. V. et al. The mTORC1/4E-BP1 axis represents a critical signaling node during fibrogenesis. Nat Commun 10, 6 (2019).

29. Weng, H.L. et al. Effect of interferon-gamma on hepatic fibrosis in chronic hepatitis B virus infection: a randomized controlled study. Clin Gastroenterol Hepatol 3, 819–828 (2005).

30. Luo, X.-Y. et al. IFN-γ deficiency attenuates hepatic inflammation and fibrosis in a steatohepatitis model induced by a methionine- and choline-deficient high-fat diet. Am J Physiol Gastrointest Liver Physiol 305, G891–G899 (2013).

31. Eckle, S. B. G. et al. A molecular basis underpinning the T cell receptor heterogeneity of mucosal-associated invariant T cells. J Exp Med 211, 1585–1600 (2014).

32. Chen, Y. & Tian, Z. Innate lymphocytes: pathogenesis and therapeutic targets of liver diseases and cancer. Cell Mol Immunol 18, 57–72 (2021).

33. Walsh, D. A. et al. The functional requirement for CD69 in establishment of resident memory CD8+ T cells varies with tissue location. J Immunol 203, 946–955 (2019).

34. Cibrián, D. & Sánchez-Madrid, F. CD69: from activation marker to metabolic gatekeeper. Eur J Immunol 47, 946–953 (2017).

35. Drescher, H. K. et al. L-Selectin/CD62L is a key driver of non-alcoholic steatohepatitis in mice and men. Cells 9, 1106 (2020).

36. Khan, O. et al. TOX transcriptionally and epigenetically programs CD8+ T cell exhaustion. Nature 571, 211–218 (2019).

37. Lei, L. et al. Th17 cells and IL-17 promote the skin and lung inflammation and fibrosis process in a bleomycin-induced murine model of systemic sclerosis. Clin Exp Rheumatol 34 Suppl 100, 14–22 (2016).

38. Fabre, T. et al. Type 3 cytokines IL-17A and IL-22 drive TGF-β-dependent liver fibrosis. Sci Immunol 3, eaar7754 (2018).

39. Hasan, S. A. et al. Role of IL-17A and neutrophils in fibrosis in experimental hypersensitivity pneumonitis. J Allergy Clin Immunol 131, 1663–1673.e5 (2013).

40. Tao, H. et al. Differential controls of MAIT cell effector polarization by mTORC1/mTORC2 via integrating cytokine and costimulatory signals. Nat Commun 12, 2029 (2021).

41. Wang, N. I., Ninkov, M. & Haeryfar, S. M. M. Classic costimulatory interactions in MAIT cell responses: from gene expression to immune regulation. Clin Exp Immunol 213, 50–66 (2023).

42. Koay, H.-F. et al. Diverse MR1-restricted T cells in mice and humans. Nat Commun 10, 2243 (2019).

43. Tenorio, E. P., Olguín, J. E., Fernández, J., Vieyra, P. & Saavedra, R. Reduction of Foxp3+ cells by depletion with the PC61 mAb induces mortality in resistant BALB/c mice infected with Toxoplasma gondii. J Biomed Biotechnol 2010, 786078 (2010).

44. Guilliams, M. et al. Spatial proteogenomics reveals distinct and evolutionarily conserved hepatic macrophage niches. Cell 185, 379–396.e38 (2022).

45. Yao, T., Shooshtari, P. & Haeryfar, S. M. M. Leveraging public single-cell and bulk transcriptomic datasets to delineate MAIT cell roles and phenotypic characteristics in human malignancies. Front Immunol 11, 1691 (2020).

46. Dusseaux, M. et al. Human MAIT cells are xenobiotic-resistant, tissue-targeted, CD161hi IL-17-secreting T cells. Blood 117, 1250–1259 (2011).

47. Jin, S. et al. Inference and analysis of cell-cell communication using CellChat. Nat Commun 12, 1088 (2021).

48. Gutierrez-Arcelus, M. et al. Lymphocyte innateness defined by transcriptional states reflects a balance between proliferation and effector functions. Nat Commun 10, 687 (2019).

49. Maslennikov, R. et al. Gut microbiota and bacterial translocation in the pathogenesis of liver fibrosis. Int J Mol Sci 24, 16502 (2023).

50. Simbrunner, B. et al. Bacterial translocation occurs early in cirrhosis and triggers a selective inflammatory response. Hepatol Int 17, 1045–1056 (2023).

51. Lett, M. J. et al. Stimulatory MAIT cell antigens reach the circulation and are efficiently metabolised and presented by human liver cells. Gut 71, 2526–2538 (2022).

52. Li, Y. et al. Mucosal-Associated Invariant T cells improve nonalcoholic fatty liver disease through regulating macrophage polarization. Front Immunol 9, 1994 (2018).

53. Lachar, J. & Bajaj, J. S. Changes in the microbiome in cirrhosis and relationship to complications: hepatic encephalopathy, spontaneous bacterial peritonitis, and sepsis. Semin Liver Dis 36, 327–330 (2016).

54. Wan, S., Nie, Y., Zhang, Y., Huang, C. & Zhu, X. Gut Microbial Dysbiosis Is Associated With Profibrotic Factors in Liver Fibrosis Mice. Front Cell Infect Microbiol 10, 18 (2020).

55. Xia, Q. et al. The phenotypic and functional characteristics of intrahepatic CD69+CD103+ tissue-resident MAIT cells in primary biliary cholangitis. J Autoimmun 154, 103442 (2025).

56. Gu, Y. et al. CD69 expression is negatively associated with T-cell immunity and predicts antiviral therapy response in chronic hepatitis B. Ann Lab Med 45, 185–198 (2025).

57. Sibbertsen, F. et al. Expansion of CD103+CD69+CD8+ cytotoxic liver tissue resident memory T cells and inflammatory monocytes in advanced biliary atresia. Front Immunol 16, 1567645 (2025).

58. Huang, B. et al. NUDT1 promotes the accumulation and longevity of CD103+ TRM cells in primary biliary cholangitis. J Hepatol 77, 1311–1324 (2022).

59. Sobkowiak, M. J. et al. Tissue-resident MAIT cell populations in human oral mucosa exhibit an activated profile and produce IL-17. Eur J Immunol 49, 133–143 (2019).

60. Gnirck, A.C. et al. Mucosal-associated invariant T cells contribute to suppression of inflammatory myeloid cells in immune-mediated kidney disease. Nat Commun 14, 7372 (2023).

61. Kartasheva-Ebertz, D. et al. IL-17A in Human Liver: Significant Source of Inflammation and Trigger of Liver Fibrosis Initiation. Int J Mol Sci 23, 9773 (2022).

62. Gu, L., Deng, W.-S., Sun, X.-F., Zhou, H. & Xu, Q. Rapamycin ameliorates CCl4-induced liver fibrosis in mice through reciprocal regulation of the Th17/Treg cell balance. Mol Med Rep 14, 1153–1161 (2016).

63. Tan, Z. et al. IL-17A plays a critical role in the pathogenesis of liver fibrosis through hepatic stellate cell activation. J Immunol 191, 1835–1844 (2013).

64. Zhang, X.-W. et al. Antagonism of Interleukin-17A ameliorates experimental hepatic fibrosis by restoring the IL-10/STAT3-suppressed autophagy in hepatocytes. Oncotarget 8, 9922–9934 (2017).

65. Ikeno, Y. et al. Foxp3+ regulatory T cells inhibit CCl4-induced liver inflammation and fibrosis by regulating tissue cellular immunity. Front Immunol 11, 584048 (2020).

66. Kurt, A. S. et al. IL-2 availability regulates the tissue specific phenotype of murine intra-hepatic Tregs. Front Immunol 13, 1040031 (2022).

67. Atif, M., Warner, S. & Oo, Y. H. Linking the gut and liver: crosstalk between regulatory T cells and mucosa-associated invariant T cells. Hepatol Int 12, 305–314 (2018).

68. Zhou, C. et al. MAIT cells confer resistance to Lenvatinib plus anti-PD1 antibodies in hepatocellular carcinoma through TNF-TNFRSF1B pathway. Clin Immunol 256, 109770 (2023).

69. Toubal, A. et al. Mucosal-associated invariant T cells promote inflammation and intestinal dysbiosis leading to metabolic dysfunction during obesity. Nat Commun 11, 3755 (2020).

70. Memarnejadian, A. et al. PD-1 blockade promotes epitope spreading in anticancer CD8+ T cell responses by preventing fratricidal death of subdominant clones to relieve immunodomination. J Immunol 199, 3348–3359 (2017).

71. Ishak, K. et al. Histological grading and staging of chronic hepatitis. J Hepatol 22, 696–699 (1995).

72. Shaler, C. R. et al. MAIT cells launch a rapid, robust and distinct hyperinflammatory response to bacterial superantigens and quickly acquire an anergic phenotype that impedes their cognate antimicrobial function: Defining a novel mechanism of superantigen-induced immunopathology and immunosuppression. PLoS Biol 15, e2001930 (2017).

73. Luo, Y. et al. Single-cell transcriptomic analysis reveals disparate effector differentiation pathways in human Treg compartment. Nat Commun 12, 3913 (2021).

